# Cortex-wide fast activation of VIP-expressing inhibitory neurons by reward and punishment

**DOI:** 10.1101/2022.04.27.489695

**Authors:** Zoltán Szadai, Hyun-Jae Pi, Quentin Chevy, Katalin Ócsai, Florin Albeanu, Balázs Chiovini, Gergely Szalay, Gergely Katona, Adam Kepecs, Balázs Rózsa

**Affiliations:** Laboratory of 3D functional network and dendritic imaging, Institute of Experimental Medicine, Budapest-1083, Hungary; Cold Spring Harbor Laboratory, Cold Spring Harbor, NY, USA; MTA-PPKE ITK-NAP B – 2p Measurement Technology Group, The Faculty of Information Technology, Pázmány Péter Catholic University, Budapest-1083, Hungary; Volen Center for Complex Systems, Biology Department, Brandeis University, Waltham, MA, USA; János Szentágothai Doctoral School of Neurosciences, Semmelweis University, Budapest, Hungary; Departments of Neuroscience and Psychiatry, Washington University School of Medicine, St. Louis, MO, USA

## Abstract

Reward and punishment powerfully inform ongoing behaviors and drive learning throughout the brain, including neocortex. Yet it remains elusive how these global signals are represented and impact local cortical computations. Previously we found that in auditory cortex, VIP-expressing interneurons are recruited by reinforcement feedback. Here, we used 3D random-access two-photon microscopy and fiber photometry to monitor VIP neural activity in dozens of cortical areas while mice learned an auditory decision task. We show that reward and punishment evoke a rapid, cortex-wide activation of most VIP interneurons. This global recruitment mode of VIP interneurons showed variations in temporal dynamics in individual neurons and across areas. Neither their weak sensory tuning in visual cortex, nor their arousal state modulation was predictive of reinforcer responses of VIP interneurons. We suggest that VIP-expressing cortical inhibitory neurons transduce global reinforcement signals to provide disinhibitory control over local circuit computations and their plasticity.

## INTRODUCTION

Neocortex can be divided into a number of functionally distinct areas such as the visual, frontal, and motor cortical regions, each specializing in different roles (Felleman and Van Essen, 1991). Classical studies have established that the specialization of each region is reflected in their neural responses; for instance, neurons in the visual cortex respond to information about the visual world, while neurons in the motor cortex inform about actions. There is an additional layer of mechanisms known to modulate these cortical responses, spanning from the broad effects of arousal to the location-specific impact of attention (Harris and Thiele, 2011). Intriguingly, there is also a growing body of evidence suggesting that each area can represent non-classical features such as reward timing (Monk et al., 2020) and category representation (Goltstein et al., 2021) in visual cortex, visual stimuli and motor modulation in the auditory cortex (Attinger et al., 2017; Nelson et al., 2013), more recently observed across other cortical regions (Allen et al., 2017; Musall et al., 2019; Stringer et al., 2019). Here we pursued a similar unexpected response pattern based on our previous observation that auditory cortex VIP interneurons respond not only to auditory stimuli but also to reward and punishment (Pi et al., 2013).

VIP expression demarcates a small interneuron subpopulation (15-20%) located mostly in the upper layers of the cortex (Acsady et al., 1996; Kim et al., 2017). Previous studies have identified a cortical circuit motif controlled by VIP interneurons that preferentially inhibit other interneurons and thereby disinhibit principal neurons (Lee et al., 2013; Pfeffer et al., 2013; Pi et al., 2013). In this circuit, VIP interneurons mainly inhibit somatostatin interneurons, which tend to exert an inhibitory drive on the dendrites of cortical pyramidal neurons (Gentet et al., 2012). Such disinhibition could lead to the selective amplification of local processing and serve the important computational functions of gating and gain modulation (Pi et al., 2013). Hence, one proposed role for VIP interneurons is to gate the integration and the plasticity of the synaptic inputs onto pyramidal neurons (Letzkus et al., 2015; Williams and Holtmaat, 2019). The same stereotyped connectivity was found in functionally and cytoarchitectonically different regions of the brain, across the auditory, prefrontal (Pi et al., 2013), visual (Pfeffer et al., 2013), and somatosensory (Gasselin et al., 2021; Lee et al., 2013) cortices, and in the amygdala (Krabbe et al., 2019). However, it is not known whether VIP interneurons have similarly stereotyped functional roles across cortical regions.

VIP interneurons have been shown to have a multiplicity of roles in sensory processing, arousal modulation, learning, and plasticity. First, studies in the primary sensory regions – barrel, auditory, and visual cortices – have demonstrated that tactile, auditory, and visual stimuli drive VIP neuron activity in diverse ways (Ibrahim et al., 2016; Khan et al., 2018; Kuchibhotla et al., 2017; Mesik et al., 2015; Pi et al., 2013; Sachidhanandam et al., 2016). However, the sensory tuning of VIP neurons tends to be weak compared to that of principal neurons. Second, VIP interneuron activity is highly correlated with the changes in pupil dilation and locomotion, suggesting a role in modulating cortical processing across arousal states (Dipoppa et al., 2018; Fu et al., 2014; Garcia-Junco-Clemente et al., 2017; Jackson et al., 2016; Pakan et al., 2016; Reimer et al., 2014; Zhang et al., 2014), while other reports show that locomotion modulates sensory processing independently from VIP activation (Yavorska and Wehr, 2021). Finally, optogenetic or pharmacogenetic inhibition (Donato et al., 2013; Fu et al., 2015; Kamigaki and Dan, 2017) of VIP interneurons, as well as their developmental dysregulation (Batista-Brito et al., 2017; Fu et al., 2015) impairs learning and plasticity in sensory discrimination and memory-guided tasks (Batista-Brito et al., 2017; Kamigaki and Dan, 2017).

We sought to investigate common rules that recruit VIP interneurons. Our starting point was the observation that auditory cortical VIP neurons respond not only to auditory stimuli but also to reward and punishment (Pi et al., 2013). VIP cells have been reported to respond to reward in hippocampus and medial prefrontal cortex and to foot shock in amygdala (Krabbe et al., 2019; Pinto and Dan, 2015; Turi et al., 2019). Those later observations fit the function of these areas in learning and plasticity. In contrast, such activity was not found in the dorsal cortex (Khan et al., 2018; Sachidhanandam et al., 2016). This questions the existence of a global reinforcement-related VIP interneuron recruitment that would support associative learning. To address this, we set up to systematically record VIP interneurons across the whole dorsal cortex during an auditory decision task. To allow simultaneous monitoring of large number of VIP interneurons across a variety of cortical regions, we used 3D acousto-optical (AO) two-photon microscopy, providing both a high signal-to-noise ratio (SNR) and high temporal resolution across large volumes. To gain access to deeper-lying cortical regions like medial prefrontal and auditory cortices, we used fiber-photometry and measured the bulk activity of VIP interneurons. We show that most VIP interneurons across cortex are indeed robustly activated by reward and/or punishment, and regional and task related behavioral factors contribute to shape their response profile differently. This global mode of recruitment of VIP interneurons is distinct from known arousal modulation of their activity and separate from the local response mode of VIP interneurons.

## RESULTS

### Auditory discrimination task for mice

To probe the behavioral function of VIP interneurons, we trained head fixed mice (n=16) on a simple auditory discrimination task (**Figure 1A**). Each trial began with the delivery of a 0.5 s auditory stimulus and mice were trained to lick (go trials) or withhold licking (no-go trials) based on the tone identity. Successful licking after tone delivery during go trials was rewarded with water (hit trials), while the absence of licking was not rewarded (miss trials). Licking for no-go trials triggered a mild air-puff punishment (false alarm, FA), which was omitted if the animal successfully withheld licking (correct rejection, CR). Mice learned this task over 3±0.6 (mean±SD) sessions after introducing the no-go tone, reaching a performance level of 80% (percentage of correct responses, Hit or CR). All recordings in this study were obtained early in training in order to also investigate VIP interneuron air puff, punishment-mediated responses (FA trials).

**Figure 1.**
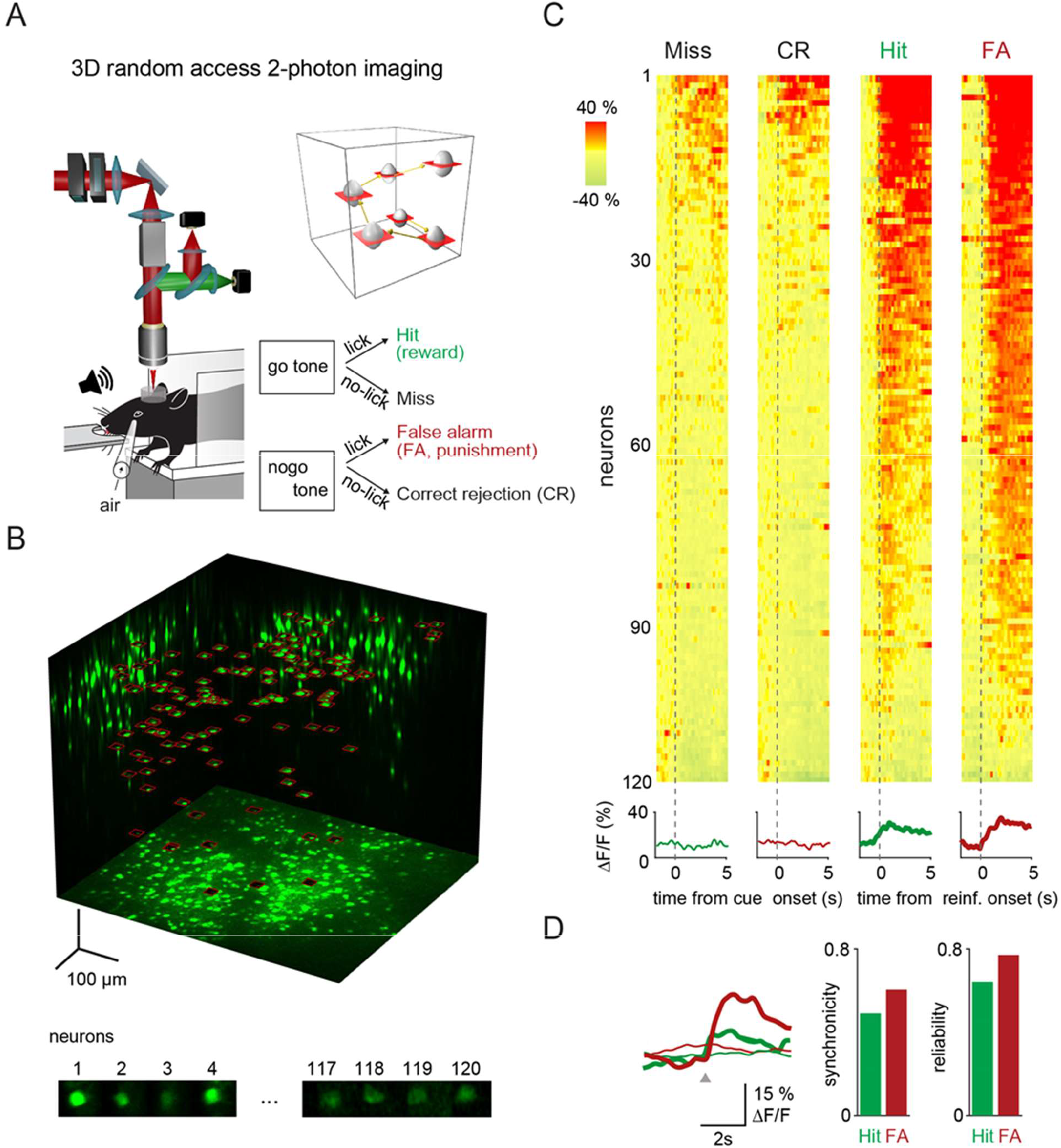
3D-random-access two-photon imaging of VIP neurons in an auditory discrimination task. **A)** Schematic of the combined fast 3D AO imaging and behavior experiments. Head-restrained mice were trained to perform a sensory discrimination, an auditory go-no-go task during 3D AO imaging using the chessboard scanning method (inset). **B)** Top, maximal intensity z and two side projections of the GCaMP6f-labeled VIP interneuron population imaged by fast 3D AO scanning. All 120 neurons within the cubature were simultaneously imaged using 120 frames of chessboard scanning (red frames). Bottom, exemplified image frames of chessboard scanning. Frames of chessboard scanning captures not only somata of the neurons but also the surrounding background information. In this way, fluorescence information is preserved during brain motion in behaving animals for motion correction. **C)** Top, somatic Ca^2+^ responses recorded during example Miss, CR, Hit, and FA trials were aligned to the reward and punishment onset, and for Miss and CR trials, to the cue onset. Responses were ordered according to their maximum amplitude. Bottom, mean±SEM responses. **D)** Left, average transients for Hit (thick green), FA (thick red), Miss (thin green) and CR (thin red) responses recorded from the 120 VIP interneurons. Right, average synchrony (mean ± SEM) and trial-to-trial repeatability (reliability) of the individual neuronal responses. Grey triangle marks the reinforcement onset in case of Hit and FA.

### Imaging VIP neurons with fast 3D acousto-optical microscopy

To study the reinforcer-mediated dynamics of VIP interneurons across the cortex, we sought to simultaneously record a large number of VIP cells, across a large cortical volume. Because of their sparse cortical distribution, electrophysiological methods combined with optogenetics-assisted identification are less suitable for cortex-wide recordings of VIP interneurons. To overcome this challenge, we used random-access, three-dimensional, acousto-optical (3D-AO), two-photon microscopy (Katona et al., 2012; Nadella et al., 2016; Szalay et al., 2016). This method allows to restrict the measurement time solely to the regions of interests. Additionally, two-photon fluorescence excitation results in high imaging penetration required for *in vivo* imaging while also delivering high spatial resolution, therefore limiting neuropil contamination (Helmchen and Denk, 2005; Horton et al., 2013; Yildirim et al., 2019). Here, we used 3D chessboard scanning (Szalay et al., 2016) that generates small patches encompassing each neuron soma. This scanning mode preserves fluorescence information during brain movements and thereby allows motion correction in behaving animals (**Figures 1A and 1C**, for theoretical summary see (Marosi et al., 2019). Overall, chessboard scanning produces an additional ~170-fold increase in measurement speed and ~15-fold increase in SNR, compared to a high-speed resonant mirror-based system scanning the same volume (**Table S1**). Thus, we could simultaneously image the activity of up to 120 GCaMP6f-expressing VIP cells (range: 12-120 cells) in a 689 μm × 639 μm × 580 μm scanning volume at a minimum of 27.8 Hz rate (**Figures 1B and 1C**).

### VIP neurons are simultaneously activated by reward and punishment in parietal cortex

We first focused on measuring the calcium-related activity of VIP interneurons in the medial parietal association area (MPta). **Figure 1** shows an example recording of 120 VIP interneurons from the MPta while the animal performed the auditory discrimination task described above. We found that the majority of VIP interneurons responded to reward and punishment presentation (reward = 85%, punishment = 90%, reward and punishment = 75% of recorded VIP interneurons). Individual neurons showed a high reliability in their recruitment (percentage of active trials for a given neuron) of 64% and 77% for Hit and FA trials, respectively. Examining individual trials, 49% and 60% of VIP interneurons were simultaneously activated by reward and punishment, respectively (**Figure 1D**). On the contrary, PV interneurons, another class of GABAergic interneurons did not show a comparable homogeneity in their recruitment by primary reinforcers. Reward and punishment delivery induced an increase in activity of respectively 29% and 10% of PV interneurons recorded in MPta **(Figure S1E).**

### VIP neurons are activated by reward and punishment across dorsal cortex

We then extended recordings of VIP interneurons to most of dorsal cortex including visual, somatosensory, motor and parietal areas (**Figure 2A**, 16 mice, one to two areas per mouse). Among the 811 neurons imaged, 65 VIP interneurons did not show statistically significant responses to behavioral events (e.g. auditory or visual stimulation, reward or punishment delivery) and were therefore excluded from further analyses. 83% and 85% of the remaining 746 VIP interneurons, responded to reward and punishment, respectively (**Figure 2D**). We found that 73% of the VIP interneurons significantly responded to both reward and punishment, similar to our observations in MPta. Further, the response of VIP interneurons to reward and punishment showed a strong correlation (Pearson correlation coefficient for average amplitudes: 0.73, **Figure 2E**). Reliable co-activation of VIP interneurons was also observed in our recordings extending throughout the dorsal cortex (**Figure S2**). On a given trial, 58% of VIP interneurons were simultaneously activated (57 ± 2.4% and 58 ± 2.5%, for Hit and FA trials respectively, **Figure S2D**) with a reliability of 61% (59 ± 1.7% and 63 ± 2.4%, for Hit and FA trials respectively, **Figure S2C**). In contrast, only 15% of the VIP interneurons responded to auditory cues in miss and correct rejection trials (**Figure 2C**).

**Figure 2.**
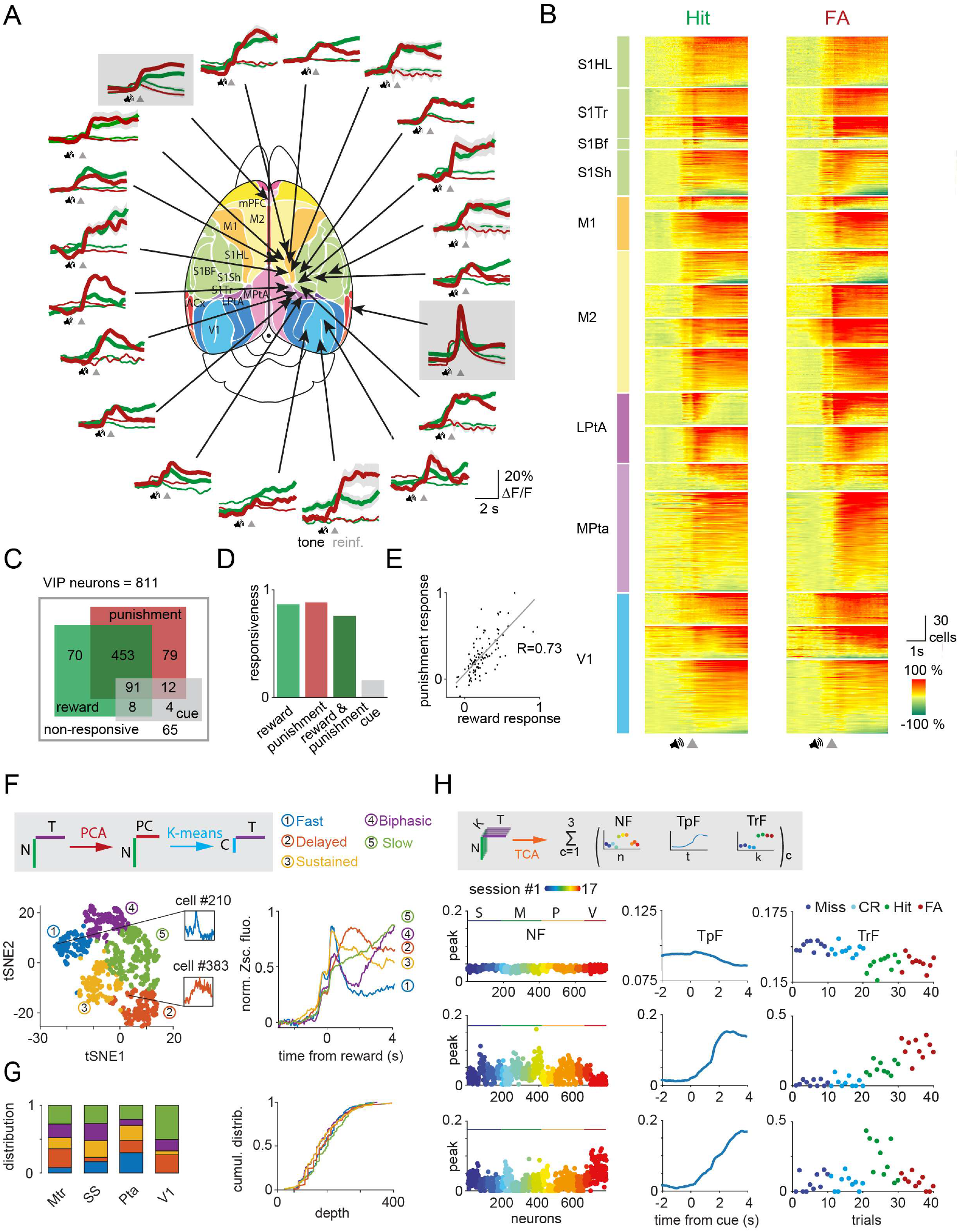
Reward and punishment recruit VIP neuronal activity across the dorsal cortex. A) Ca^2+^ responses of individual VIP interneurons recorded separately from 18 different cortical regions with fast 3D AO imaging were averaged for Hit (thick green), FA (thick red), Miss (thin green), CR (thin red). Fiber photometry data were recorded simultaneously from mPFC and ACx regions and are shown in gray boxes. Functional map (Kirkcaldie, 2012) used with the permission of the author. B) Each line of the raster plots shows average neuronal response for Hit and FA. Abbreviations indicate color coded cortical recording positions shown in panel A. Responses were normalized in each region and ordered according to their maximum amplitude. **C)** Responsiveness of 811 VIP interneurons for Hit and FA. **D)** Bar chart of data from C. **E)** Average response of individual VIP interneurons for FA as a function of the response for Hit. Note the high correlation (R=0.73). **F)** Left, T-distributed Stochastic Neighbor Embedding (tSNE) plot of the reward mediated activity of VIP interneurons after PCA. Individual neurons are color coded according to their cluster type obtained using a k–means clustering algorithm. Inserts show the average response of single rapidly (blue) or delayed (orange) activated VIP interneurons. Right, average GCaMP6f responses from different clusters of VIP interneurons after reward delivery. **G)** Left, distribution of the clusters shown in panel F across different cortical areas. Right, cumulative distribution of the clusters shown as a function of cortical depth. **H)** Top, schematics of temporal component analysis. Below: rank 3 TCA neural, temporal, and trial factors. Miss and CR trial factors were indicated here with dark and light blue dots. The second component clearly distinguishes between trials with and without reinforcement.

### Recruitment of VIP interneuron population by reward and punishment in the medial prefrontal and auditory cortex

To probe additional but deep-lying cortical structures, we took advantage of the coherent recruitment of VIP interneurons by reinforcers and used fiber photometry (Cui et al., 2013; Gunaydin et al., 2014). This approach allowed us to simultaneously measure bulk calcium-dependent signals from VIP interneurons located in the right medial prefrontal (mPFC) and left auditory cortices (ACx) by implanting two 400 μm optical fibers at these locations (n=6 mice, **Figure S1C**). Consistent with our previous electrophysiological results in ACx (Pi et al., 2013) and two-photon imaging from dorsal cortical regions, calcium-related signals from VIP interneurons in the ACx and mPFC were increased after reward and punishment delivery (in ACx: Hit = 4.8 ± 0.32 %, FA = 10.9 ± 0.03 %; in mPFC: Hit = 4.3 ± 0.69 %, FA = 6.6 ± 0.85 %, ΔF/F peaks, **Figure 2A**). We did not further analyze the FA responses in auditory cortex as those responses also had a sensory component linked to the white noise-like sound created by the air puff delivery. Similar to our single cell results, PV-expressing neuronal population in ACx did not show any significant change in activity after random reward delivery (**Figure S2F**). Concurrent recordings of VIP interneuron population in ACx and mPFC revealed heterogeneity in the dynamics of VIP interneuron activity during reward delivery (**Figure 2A**). VIP interneurons in the auditory cortex showed a phasic-like response to reward (peak time for Hit = 0.06 ± 0.036 s, decay time constant = 2.7 s). In contrast, medial prefrontal VIP interneurons were slowly activated (peak time for Hit = 3.08 ± 0.968 s, decay time constant = 7.75 s, **Figure 2A**). These population recordings confirmed the dominant contribution of reinforcement-related signals to VIP interneuron population responses but also reveal potential area-specific heterogeneity in the dynamics of VIP interneuron activity.

### Heterogeneity in the dynamics of reinforcer-related activity of individual VIP interneuron

The difference in dynamics at the population level across different brain areas might be supported by heterogeneity in the individual response profiles of VIP interneurons. Thus, we sought to characterize the dynamics of VIP interneurons at a single cell resolution and across dorsal cortex using chessboard 3D AO recordings. We first focused on VIP interneurons activated upon reward delivery during the sensory discrimination task (n=606 cells). We applied principal component analysis (PCA) to the average reward responses of individual neurons to reward. We then clustered these responses using k-means clustering. This approach did not primarily separate neurons according to the recording sessions (**Figure S3A**). Rather, our clustering approach allowed us to delineate 5 groups of VIP interneurons (**Figure 2F**). Based on visual inspection of their mean temporal profiles, we labeled these groups as: ‘fast’ (n=109), ‘delayed’ (n=88), ‘sustained’ (n=177), ‘biphasic’ (n=120) and ‘slow’ (n=112). Note that all of these response types share important similarities such as a phasic reward response and mostly differ in their subsequent temporal dynamics. We first considered the distribution of these 5 types of neuronal responses across different brain areas. We observed an overrepresentation of the ‘fast’ group in parietal cortex and of the ‘slow’ group in primary visual cortex (**Figure 2G**). The ‘fast’ group was absent from visual cortex (**Figure 2G**). To quantify this heterogeneity across cortical areas, we defined 5 feature vectors as the mean response of each cluster to rewards and then projected the reward response of each VIP interneuron onto these features (**Figure S3B**). We found that the projection associated with the ‘fast’ group were significantly higher for VIP interneurons located in parietal compared to those recorded in visual cortex (mean Δ_Pta-V1_=3.22, Mann-Whitney test, p=3.77 10^-9^), while the opposite was observed for the projection associated with the ‘slow’ group (mean Δ_Pta-V1_=−7.79, p=3.77 10^-9^**, Figure S3B**). Finally, we took advantage of the 3D AO imaging to investigate the heterogeneity in the responses of VIP interneurons located at different depth of the cortex. We were able to detect some differences in the amplitude of the average responses for reward (**Figure S2E**, F= 9.5, p=0.002). However, we did not observe any differences in the distribution of the different clusters across depth (**Figure 2G**, F=1.16, p=0.36).

The differences in average response dynamics from individual neurons could arise from inter-trial variability. To evaluate this potential heterogeneity in the single trial dynamics of VIP interneuron activity, we used tensor component analysis (TCA, **Figure 2H**). TCA allowed us to further characterize the trial identity-dependent dynamics of VIP interneuron activity. All recorded neurons from different sessions/cortical regions were grouped by keeping only the first 10 trials of each trial types (see Methods for additional information) We used nonnegative tensor decomposition and focused our analysis on rank 3 TCA as using higher rank showed signs of overfitting and did not improve the reconstruction error (22% for rank 3 vs 18% for rank 20). We found a latent temporal component that robustly separated hit and false alarm from miss and correct rejection trials (2^nd^ component, **Figure 2H**, mean_miss&CR_ vs mean_hit&FA_ = 0.02 vs 0.2, Mann-Whitney test, p<0.001). The third latent temporal component showed a slower time course, with some reward specificity and was over-represented in neurons from the visual cortex (mean_SS,Mtr,Pta_ vs mean_V1_ = 0.02 vs 0.06 p<0.001).

### Behavioral performance influences task-related VIP interneuron responses

Differences in individual animal performance of the discrimination task could also contribute to the heterogeneity in the activity of VIP interneurons. Hence we tested whether differences in hit rate influenced the response of VIP interneurons during various epochs. We observed a positive correlation between the hit rate and the magnitude of the cue response of VIP interneurons during hit trials (R=0.62, **Figure S3D**). Using a simple linear regression model, we found that the hit rate was able to explain 39% of the variance of cue responses (R^2^=39.0% p=0.006). For comparison, the cue response was not influenced by the location of VIP interneurons in the dorsal cortex (R^2^ = 18.2% p=0.41).

### Arousal modulation of reinforcement-mediated recruitment of VIP interneurons

Reward and punishment can induce changes in the arousal states of the animals, and the activity of VIP interneurons is known to be modulated by the arousal states (Fu et al., 2014; Garcia-Junco-Clemente et al., 2017; Reimer et al., 2014). Therefore we considered whether changes in arousal contributed to the recruitment of VIP interneurons by primary reinforcers. We monitored variations in pupil diameter as a proxy for assessing arousal states (Vinck et al., 2015). In this set of experiments, we restricted our measurements to somatosensory and motor cortices using 3D AO microscopy and to the auditory and the medial prefrontal cortex using fiber photometry as described above (**Figure 3A**).

**Figure 3.**
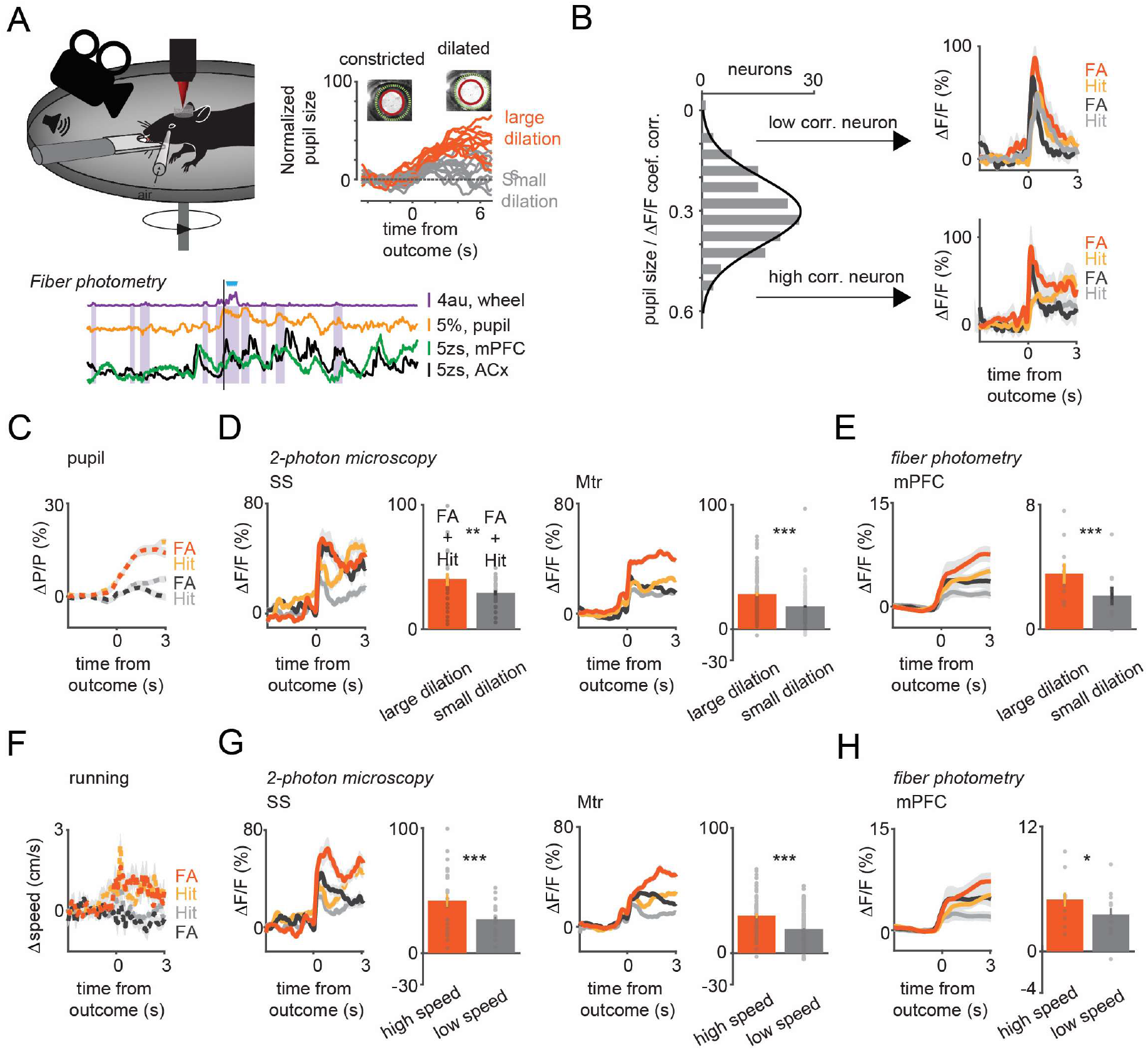
Arousal states modulate VIP neural responses to sensory cues and reinforcers. A) Upper left, schematic of measurements. Pupil and movement were simultaneously monitored during 3D imaging in the auditory go-no-go task. Upper right, high (orange) and low (gray) arousal states were separated by changes in pupil diameter. Below, 60 sec continuous monitoring of different behavioral variables together with VIP interneuron population activity in ACx and mPFC. The black bar indicates the timing of an uncued reward delivery. Blue triangles indicate licking events. Purple shaded boxes represent running bouts. B) Left, distribution of correlation coefficients of relative change in pupil diameter (ΔP/P) and VIP neuronal response. Right, reinforcement-associated responses were significantly larger when relative change in pupil diameter (ΔP/P) was higher during the task. Red and orange indicate FA and Hit responses associated with higher ΔP/P. FA and Hit responses associated with low ΔP/P are in black and gray, respectively. **C)** Average pupil dilation traces during high (red and orange) and low (black and gray) pupil changes for FA and Hit trials. **D)** Population averages for Hit and FA during high and low pupil change in the SS and Mtr regions. Bars indicate average peak amplitudes (mean±SEM, Hit and FA combined). Even in the late period, when the outcome responses were dissipated, larger changes in pupil diameter at the time of reinforcement were associated with higher VIP responses. **E)** Same as D but for fiber photometry in the mPFC. **F)** Same as C but for running speed. **G)** Same as D but for running speed. **H)** Same as E but for running speed. Higher relative change in the running speed was associated with larger neuronal responses recorded with 3D imaging or fiber photometry.

Hit and false alarm trials were first split into two groups using the mean reinforcer-mediated pupil dilation for threshold (average changes in pupil size for the large and small pupil group:14.38% vs. 2.81%; **Figure 3C**). Reinforcement delivery associated with bigger changes in pupil diameter led to a stronger recruitment of VIP interneurons in both the somatosensory (large vs. small pupil ΔF/F: 40% vs 29% n=26, t-test, p=0.01) and motor cortex (large vs. small pupil ΔF/F: 28% vs 18%, n=111, P<0.001 **Figure 3D**). A comparable modulation of VIP interneuron activity upon reinforcer presentation was observed when trials were split based on baseline pupil diameter (see Supplemental Information and **Figure S4B**). The recruitment of VIP interneurons upon cue presentation only (i.e. Miss and CR trials, where trials were split using the mean cue-mediated pupil dilation for threshold) was similarly modulated by arousal (**Figure S4A**). The positive correlation between pupil size changes and reinforcer-related activity of VIP interneurons was also present at single cell level (median correlation coefficient 0.31). Interestingly, neurons with a strong correlation coefficient showed a slower activity profile of recruitment by reinforcers than those with a correlation coefficient below the median value (**Figure 3B**)

Because VIP interneuron population had slower dynamics in medial prefrontal cortex than in auditory cortex, we hypothesize that the pupil size-dependent modulation of reward responses would be stronger in prefrontal cortex than in auditory cortex. Reinforcer-mediated responses in medial prefrontal cortex were significantly larger in trials with greater changes in pupil diameter (large vs. small pupil ΔF/F: 3.8% vs 1.4%, n=6 mice, *p*=0.01 **Figures 3E and S4C**). This pupil size dependent modulation was however absent in recordings of VIP interneurons in the ACx (large vs. small pupil ΔF/F: 4.5% vs 3.2%, n=6 mice, *p*=0.08 in the ACx (**Figure S4A**). Similarly, to our single neuron measurements, arousal modulation was present at a population level in Miss and CR trials (**Figure S4A**) or when using the baseline pupil dilation as arousal index (**Figure S4B**).

### Modulation of reinforcement-mediated recruitment of VIP interneurons by locomotion

We next examined how running behavior modulates VIP activity. Response profiles were split based on the median speed change during reward and air puff delivery (average speed for the fast and slow group 1.20 cm/s vs. −0.23 cm/s **Figure 3F**). Similar to what we observed with the pupil size, VIP interneuron responses to reward were stronger when the mice ran faster both for somatosensory (fast vs. slow running ΔF/F: 42% vs 27% n=26, t-test, *p*<0.001) and motor cortex-located interneurons (fast vs. slow running ΔF/F 30% vs 19%, n=86, *p*<0.001, **Figure 3G**). This difference was also found for sensory responses during Miss and CR (**Figure S4D**). Similar to the results of pupil modulation, we also observed that VIP interneuron population shows a stronger modulation by speed change in mPFC compared to ACx (fast vs. slow running mPFC, ΔF/F: 5% vs 3.5%, *p*=0.02, ACx, ΔF/F: 5.4% vs 4.1%, p=0.44, **Figures 3H and S4D**).

### VIP neurons in visual cortex are activated by sensory stimuli and reinforcers

VIP interneurons in auditory and visual cortices respond to sensory stimulation, albeit their responses are weak tuned (Kerlin et al., 2010; Mesik et al., 2015; Pi et al., 2013). Therefore we next compared sensory- and reinforcement-evoked activity of VIP interneurons in visual cortex. In this set of experiments to ensure stimulus control we lightly anesthetized mice with isoflurane and imaged responses to drifting grating bars with different orientations. The vast majority of VIP interneurons (93.5%) responded to visual stimuli. We computed the orientation selectivity index (OSI) and direction selectivity index (DSI) of each VIP neuron (see methods). As previously reported (Kerlin et al., 2010; Mesik et al., 2015), VIP interneurons showed broad tuning with little or no preferred directions or orientation (OSI = 0.17 ± 0.01, DSI = 0.16 ± 0.01, **Figures 4C**) especially compared to pyramidal cells (OSI = 0.63 ± 0.01, t-test, p<0.001, DSI = 0.34 ± 0.01, *p*<0.001).

**Figure 4.**
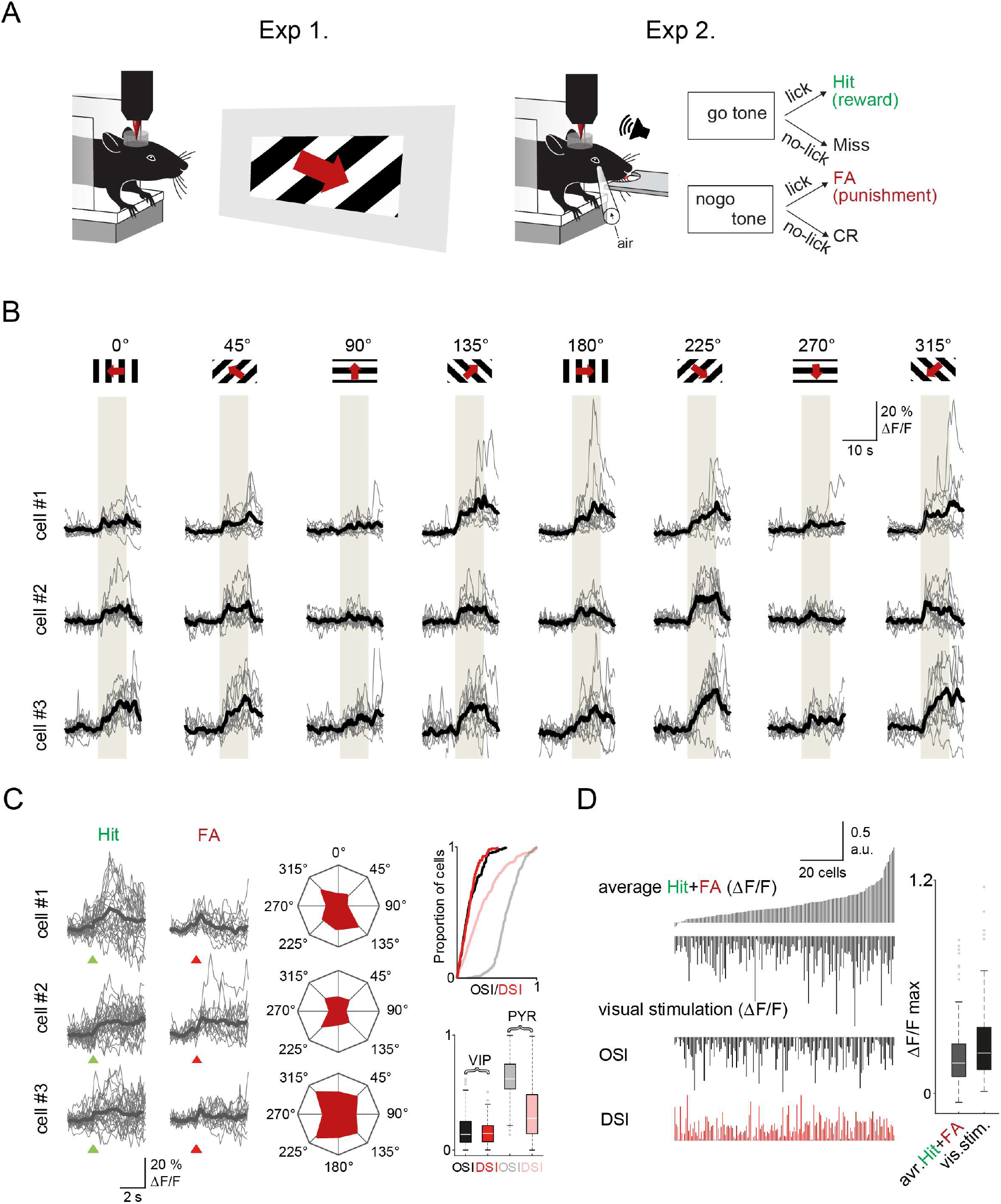
Visual cortex VIP neurons respond to both visual stimuli and reinforcers. **A)** Schematic of the measurement. Orientation tuning was mapped in a first set of experiments (Exp. 1) which was followed by recordings of the same neurons during the auditory go-no-go task (Exp. 2). Both set of recordings were performed using fast 3D AO imaging. **B)** Individual Ca^2+^ responses from three different VIP interneurons to visual stimulation with moving grating in 8 different directions. The grey boxes indicate the duration of the visual stimulation. **C)** Left, responses of the same three cells to reinforcement. Middle, polar plots of neuronal responses to visual stimulation from the same neurons. Right top, cumulative distribution plot of OSI and DSI parameters of VIP (black and red) and pyramidal cells (gray and pastel red (VIP: n=157 cells, n=3 mice, pyramidal cells: n=383 cells, n=3 mice. Right bottom, OSI and DSI values of the same cells. Box-and-whisker plots show the median, 25th and 75th percentiles, range of nonoutliers and outliers. **D)** Correlation between reinforcement and visual responses in the same VIP interneurons (n= 157). Each column refers to a single cell. From top to bottom: mean of the average Hit and FA responses, average visual responses, mean orientation selectivity index (OSI) and mean direction selectivity index (DSI). The cells were ordered according to the amplitude of the averaged reinforcement signal. Right, maximums of reinforcement-related and visual stimulation responses. Box-and-whisker plots show the median, 25th and 75th percentiles, range of nonoutliers and outliers.

After measuring their visual response tuning, we imaged the same visual cortical neurons while mice performed the auditory discrimination task (**Figure 4A**). We found that 80.4% of VIP neurons were significantly activated by reward or punishment with a response magnitude comparable to their visual responses. However, the reinforcement responses were only weakly correlated with visual stimulus-evoked responses (Pearson’s R value for reward: 0.16, for punishment: 0.23, and reward and punishment combined: 0.22; **Figures 4D and S5A**). Similarly, neither the orientation nor the direction selectivity index of the VIP neurons correlated with their reinforcement responses (Pearson’s R value for OSI: 0.08, for DSI: 0.06; **Figure S5B**). This supports the hypothesis that the global recruitment of VIP interneurons by reinforcers arises independently of the local computation these neurons might be involved in.

## DISCUSSION

Here we examined the rules of behavioral recruitment for VIP interneurons across cortex. By monitoring VIP neural activity across dozens of cortical regions we found that most neurons were strongly activated by water reward and air-puff punishment. This recruitment was boosted during high arousal states as previously observed for other sensory-mediated processes (Fu et al., 2014; Garcia-Junco-Clemente et al., 2017; Reimer et al., 2014). In visual cortex VIP interneurons also showed sensory tuning to visual gratings. This tuning was however not predictive of the VIP interneuron responses to reinforcers demonstrating the co-existence of both local and global modes for the recruitment of VIP interneurons.

VIP interneurons represent a small and sparsely distributed population across cortex rendering their investigation challenging. We aimed at simultaneously monitor, in behaving mice, the activity of a large population of these neurons and this across dorsal cortex. This was made possible by the use of 3D acousto-optical two-photon microscopy. The chessboard scanning method of 3D AO microscopy provided additional advantages to our ability to measure the spatial and temporal dynamics of VIP interneuron activity. According to our calculation this method leads to a several orders of magnitude increase in the measurement speed and signal to noise ratio compared to piezo-based volume scanning (**Table S1**). Further, it enabled robust off-line motion correction during behavioral experiments owing to the ability to actively extend the recordings beyond the soma of the neurons, thereby preserving fluorescence information during motion (Reid et al., 2016). Due to this large improvement in the SNR and recording speed (**Table S1**), we were able to dramatically increase the number of simultaneously recorded neurons while maintaining a high sensitivity of detection of neuronal activity. This allowed us to demonstrate that VIP interneurons, throughout cortex and across cortical layers, responded homogenously and synchronously to reward and punishment delivery.

This finding seemingly contradicts previous reports of muted VIP interneurons reinforcement responses in similar goal directed tasks (Khan et al., 2018, Sachidhanandam et al., 2016). Those studies however used over trained animals for which little to no punishment was delivered and reward delivery was fully predictable. One study limited to the amygdala indeed showed that reinforcement recruits VIP neurons in a time limited manner (Krabbe et al 2019).

Our observations also revealed heterogeneity in VIP interneurons: their temporal response profiles to reinforcers, sensory tuning and arousal modulation showed differences. In addition to the single trial and individual neuronal variability in the dynamics of reinforcer related activity revealed by principal and tensor component analyses, we identified variability in behavioral performance as a source of heterogeneity in the cue-mediated recruitment of VIP interneurons **Figure S3D**). We also investigated potential response heterogeneity across cortical regions in the reinforcement-mediated response of VIP interneurons. For instance, VIP interneuron population showed a faster recruitment by reward in ACx than their counterpart in mPFC (**Figures 2A, 3E and S4A**). Rapidly activated neurons were absent from visual cortical area whereas they could be observed throughout the rest of the dorsal cortex (**Figure 2G**). This variability might partially reflect different VIP interneuron subtypes (Pfeffer et al., 2013; Pi et al., 2013; Pronneke et al., 2015). Perhaps the most distinct subclass of VIP interneurons is cholecystokinin-(CCK+) expressing interneurons with basket cell-type morphology (He et al., 2016). These cells provide somatic inhibition in the hippocampus. We expect that the majority of the neurons we recorded in the upper layers are calretinin-(CR+) expressing bipolar cells, including intrinsic cholinergic acetyltransferase-(ChAT+) positive neurons (Kim et al., 2017). This separation into CCK-, CR- or ChAT-expressing VIP interneurons has been recently partially validated using single-cell transcriptomic analysis (Tasic et al., 2016; Zeisel et al., 2018). Given the high proportion of VIP neurons responding to reward and punishment, it seems likely that multiple subtypes of VIP interneurons respond to reinforcers. Further studies using inter-sectional targeting strategies will be required to provide insight into the potential cell-type-specific origins response of heterogeneity we observed.

The response heterogeneity of the local mode of VIP interneurons had already been appreciated for sensory stimuli. When local activation is examined in terms of tuning, VIP interneurons are significantly more heterogeneous and broadly tuned than principal neurons, as previously shown in the auditory and visual cortices (Mesik et al., 2015; Pi et al., 2013). Indeed, we found that VIP interneurons respond with a low selectivity for drifting grating visual stimuli. There was only a weak positive correlation between the reinforcement-related and the visual stimulus-driven responses. This small correlation could reflect differences in excitability, but more importantly indicates that VIPs can play in both leagues: in region-specific sensory processing and in transmitting global signals to local microcircuits. This further suggests the absence of distinct populations specializing only in global or in local processing.

We also observed arousal-modulation of VIP interneuron activity in motor, somatosensory and medial prefrontal cortices, consistent with the previous reports in visual and pre-motor areas (Fu et al., 2014; Garcia-Junco-Clemente et al., 2017; Jackson et al., 2016; Reimer et al., 2014). Arousal states are usually measured by changes in pupil diameter or running speed. One caveat of comparing reinforcement-evoked responses to arousal modulation is that the delivery of water reward and air puff punishment also drives additional changes in arousal, leading to pupil dilation and/or locomotion. Nevertheless, we found that VIP interneuron recruitment by reinforcers was correlated with pupil dilation, similar to the previously documented arousal-modulation (Fu et al., 2014; Garcia-Junco-Clemente et al., 2017; Jackson et al., 2016; Reimer et al., 2014). However, only the late response phase showed a correlation with the pupil size, whereas the initial, transient phase followed more closely the reinforcer delivery. This arousal modulation was surprisingly muted in auditory cortex during reward delivery. This could thereby explain the striking different kinetics observed in simultaneous mPFC and ACx photometry measurements. Additionally, our behavioral paradigm did not encourage mice to run and their small movements produced only weak modulation in VIP activity (**Figure 3**). Overall, these observations lead us to conclude that changes in arousal alone cannot explain the recruitment of VIP interneurons upon reward or punishment.

The circuit basis of the global signal is not yet known, although neuromodulatory systems are prime candidates, in particular, the forebrain cholinergic system. Indeed, central cholinergic neurons convey rapid reinforcement responses to cortex (Hangya et al., 2015) and a type of layer 1 inhibitory neuron is activated by punishment through a nicotinic mechanism (Letzkus et al., 2011). VIP neurons express fast, ionotropic nicotinic receptors and can be activated by acetylcholine in vitro (Alitto and Dan, 2012; Chen et al., 2015). Optogenetic cholinergic stimulation can also depolarize the membrane potential of VIP neurons in vivo (Gasselin et al., 2021). However, it remains unclear how this putative reinforcer-mediated cholinergic signaling would be ultimately integrated within cortex as multiple inhibitory neurons types other than VIP interneurons are known to respond to acetylcholine as well (Kuchibhotla et al., 2017). Another possibility is that the serotonergic system could convey this reinforcement signals (Cohen et al., 2015). Indeed, many (but not all) VIP neurons express 5HT3A, the ionotropic serotonin receptor and could thereby be a specific recipient for such information (Tasic et al., 2016; Zeisel et al., 2018).

At a functional level, reinforcer-induced activation of VIP interneurons is likely to produce disinhibition (Lee et al., 2013; Pi et al., 2013) and thereby gain modulation (Pi et al., 2013) through changing the balance of inhibition across the somato-dendritic axis (Pfeffer et al., 2013). This could then represent a circuit-level explanation for the broad recruitment of dendrites observed during reinforcement (Lacefield et al., 2019). At the dendritic level, disinhibition is known to favor dendritic spikes that will boost neural responses (Larkum et al., 2009; Lavzin et al., 2012; Palmer et al., 2014; Smith et al., 2013). This would support the role of VIP neurons in circuit plasticity in visual cortex (Fu et al., 2015) and hippocampus (Donato et al., 2013). Hence, VIP interneurons-mediated disinhibition provides a compelling basis for how global reinforcers could impact local cortical computations and drive learning.

In summary, the global activation mode of VIP cells could serve to gate dendritic plasticity cortex-wide and potentially associate distant neuronal assemblies that are active at the moment of reinforcement to link information about the internal and external worlds. In other words, the VIP-mediated feedback signaling may provide the required global learning signal to strengthen the reinforcement-related functional connectivity of cortical representations. In this way, VIP neurons may transiently boost the gain on learning, similar to the different phases of learning in deep networks in artificial intelligences (Guerguiev et al., 2017).

## MATERIALS and METHODS

### Animals

All experimental procedures were carried out following the guidelines of the Animal Care and Experimentation Committee of the Institute of Experimental Medicine of the Hungarian Academy of Sciences, and the Cold Spring Harbor Laboratory Institutional Animal Care and Use Committee, in accordance with the Hungarian, EU, and National Institutes of Health regulations. We used male and female adult (6-24-week-old) VIP-Cre, PV-Cre and Thy-1-Cre mice housed in small groups of 2-4 under controlled temperature and humidity conditions. They were kept on a reverse light cycle, and during the training and the experimental period the water consumption of the VIP-Cre mice was restricted to 1 ml/day after recovering from surgery. The mice had ad libitum access to food.

### In vivo imaging

Animals were anesthetized using a cocktail of fentanyl, midazolam, and meditomidine (0.05 mg, 5 mg, and 0.5 mg/kg, respectively). Ropivacaine 0.2 % was administered subcutaneously over the skull prior to the incision. After removing the skin over the top of the skull, which was then thoroughly cleaned and dried, a round craniotomy was performed using a 3 mm diameter biopsy punch and a dental drill. After the bleeding had been stopped, a double coverslip was implanted over the cranial window and fixed using a mixture of cyanoacrilate glue (Loctite Superbond) and luting cement (3M ESPE RelyX). Finally, a metal headbar was cemented to the skull using dental cement (C&B Superbond). The 3 mm diameter cranial window was positioned according to two main aspects. We centered the craniotomy on the injection site, except in motor and visual areas, where this would have resulted in transecting the sutures, which would have caused larger motion artefacts and severe bleeding from venous sinuses. During the procedure, the mice were head-fixed and laid on a heating pad to maintain stable body temperature. After the operation, the mice were woken up using a mixture containing nexodal, revertor, and flumazenil (1.2 mg, 2.5 mg and 2.5 mg/kg body weight, respectively), and put on another heating pad where they stayed until recovered enough to be finally put back in their home cages. Post-operative care consisted of a daily intraperitoneal carprofen injection (0.5 mg/ml, 500 μl) for up to 5 days, and subcutaneous Ringer lactate injection (0.1-0.15 ml) to prevent dehydration. The cranial window implantation was usually performed 2 weeks after the virus injection. Injection sites of the 18 dorsal cortical regions from 16 mice were defined on the basis of coordinates from the Allen and Paxinos brain atlases (**Figure 2A**). In the visual cortex, the correct location was further confirmed by recording the responses of the cells to visual stimulation. Post hoc histology was performed in early experiments to ensure our bregma coordinates matched the Paxinos atlas. Each region was then recorded one time per animal.

### Viral injection

Anaesthesia and post-operative care were executed as above. A small, approximately 0.5 mm diameter craniotomy was performed with a dental drill. 200-300 nl AAV9.Syn.Flex.GCaMP6f.WPRE.SV40 (Penn Vector Core) was injected using a borosilicate pipette at 350 μm depth into different cortical areas for two-photon imaging. The speed of the injection was 15-20 nl/s, and there was a 10 minute period between the end of the injection and the removal of the pipette to prevent leakage. We injected one to two areas per animal.

### Optical fiber implantation

Animals were anesthetized using isofluorane (1l/min O_2_ – 0.8% isoflurane) and placed in a stereotaxic apparatus. A small craniotomy was performed using a dental drill above the left ACx (2.50 mm posterior to the bregma and 4 mm lateral to the midline) and the right mPFC (1.75 mm anterior to bregma and 0.5 mm lateral to midline). 200 nl of AAV9.Syn.Flex.GCaMP6f.WPRE.SV40 (Penn Vector Core) was then injected at a rate of 50nl/min into the ACx (1.2 mm deep) and in the mPFC (1.5 mm deep). The fiber optic cannulas (400 μm, 0.48NA, Doric lenses) were inserted 0.4 mm above the injection sites for both locations and sealed in place using Metabond, Vitrebond and dental acrylic. Behavioral training and physiological recordings were started at least 2 weeks after surgery to allow mice to recover and the fiber to clear.

### Data collection using fiber photometry

Fiber photometry data were collected and analyzed using a custom-made photometry set up and Matlab-based software. We used a 470nm LED source (M470F3, Thorlabs) coupled to an optic fiber (M75L01) and collimation lens (F240FC-A) for GCaMP6f excitation. The 470 nm excitation light was delivered to the cannula implanted on the head animal using a second collimation lens (F240FC-A) coupled to a 400 μm, high NA, low autofluorescence optic fiber (FP400URT, custom made, Thorlabs). The emission light was collected using the same optic fiber and directed to a Newport 2151 photoreceiver using a focusing lens (ACL2541U-A, Thorlabs). Excitation (ET470/24M) and emission (ET525/50) filters, and a dichroic mirror (T495LPXR) were from Chroma Technology. The 470nm excitation light was amplitude-modulated at a frequency of 211 Hz, with a max power of 40uW, using an LED driver (LEDD1B) controlled through a National Instrument DAQ (NI USB-6341). The modulated data acquired from the photoreceiver were decoded as in Lerner et al., 2015 using a custom Matlab function (available at https://github.com/QuentinNeuro/Bpod-FunctionQC).

### Auditory discrimination task

Mice were kept on a limited access water schedule for behavioral experiments. They had to lick when they heard a go tone (frequency: 5 kHz complex tones for sessions with two-photon recordings, 3kHz pure tones for sessions with photometry recordings, duration: 0.5 s) to get small water droplets (5 μl) as a reward, and avoid licking after hearing a no-go tone (frequency: 0.5 kHz complex tones for sessions with two-photon recordings, 20kHz pure tones for sessions with photometry recordings, duration: 0.5 s) which was associated with a 100 ms-long mild air puff aimed into the eye. Reinforcement came 0.5 s after it was triggered. The intensity of the air puff was set to yield a blink response. In some experiments, we introduced two additional stimuli that were less easy to discriminate (8 kHz for go and 10 kHz for no-go tones). The addition of these cues did not reveal any significant differences in GCaMP6f signals in VIP neurons, therefore these trial types were combined as go and no-go stimuli for further analysis. Licking was detected using a custom-made infrared sensor. Behavioral data were acquired using a Bpod device, and the tones were generated using a PulsePal device (Sanders and Kepecs, 2014) and Logitech speakers. In one set of experiments we measured how pupil dilation changed during behavior. A 4× objective was attached to a CMOS camera (Basler puA 1600-60 μm) to record pupil diameter and eye movements. In another set of experiments we recorded running speed: mice were head-fixed over a rotating plastic plate allowing them to run freely. The rotation speed of the dial was recorded by an optical mouse (Urage reaper 3090, Hama) mounted upside down on the lower side of the plate.

### 3D AO microscopy

The improved microscope is designed and constructed based on the previous system reported earlier (see Figure S1 in (Szalay et al., 2016)). Briefly, short pulses were delivered by a femtosecond laser (Mai Tai, Spectra Physics). The coherent backreflection was eliminated by a Faraday isolator (BB9-5I, EOT). Thermal drift errors of optical elements were compensated for by an automatic beam-stabilization unit (BeamStab, Femtonics). The temporal dispersion was compensated for by a motorized four-prism sequence that could be automatically tuned in the 720-1100 nm wavelength range to provide the required large, negative, second-(up to 100,000 fs^2^) and third-order (up to 45,000 fs^3^) dispersion compensation (4DBCU, Femtonics). The 4DBCU unit was fine-tuned to provide the best image contrast and SNR at each wavelength in the depth. The first two water-cooled AO deflectors were filled with chirped acoustic waves whose frequencies form two orthogonal electric cylinder lenses (AO z-focusing unit). The second group of AO deflectors, with 15 mm clear optical aperture (Gooch and Housego), did the majority of lateral scanning and also compensated for the longitudinal and lateral drift of the focal spot in cooperation with the first two deflectors according to equations S1-S70 published earlier (Reid et al., 2016). These two groups of deflectors were coupled together by a telecentric relay system (using two achromat lenses, #47-318, Edmund Optics) which contained a half wave plate (AHWP10M-980, Thorlabs) to set the optimal polarization for maximal diffraction efficiency. There is a one-to-one relationship (a bijection) between non-linear radiofrequency signals and the position, speed, and direction of the moving focus spot (see Equations S1-S70 and Table S1 in (Szalay et al., 2016)). We used these quadratic questions to change the frequency of the sine wave drive to generate multiple 3D drifts from any arbitrary position at any desired speed. In this way, multiple small frames were generated (3D chessboard scanning) around each VIP cell from 10-25 lines. Therefore, not only the somatic signal but also the surrounding background information was detected: this enabled the somatic fluorescent signals to be preserved, even during brain movement, for off-line motion artefact compensation. A second telecentric relay system consisting of two achromat lenses (#47-319, Edmund Optics, G322246525, Linos) focused the diffracted light beams onto the back aperture of the objective. The back-reflected fluorescence signal was separated from the excitation beam by a long-pass dichroic with a cut-on wavelength of 700 nm (700dcrxu, Chroma Technology). Red and green channels were split using a long-pass dichroic at 600 nm (t600lpxr, Chroma Technology). The absorption filters for green and red fluorescence was centered to 520 ±30 nm and 650±50 nm, respectively (ET520/60m, ET650/100m, Chroma Technology). Two extra infrared filters (ET700sp-2p8, Chroma Technology) blocked the back-scattered excitation beam from the GaAsP photomultiplier (H10770PA-40, Hamamatsu). The entire detector assembly was fixed to the objective and moved together during setting the nominal focal plane for the 3D AO imaging to minimize the detection pathway and maximize photon collection efficiency. A 20× objective (XLUMPlanFI20×/1.0, water immersion, Olympus) with a 1.0 numerical aperture was used.

### Recording sparsely-labelled networks in 3D with AO scanning

The main advantage of 3D AO microscopy is that the entire measurement time can be restricted to the required ROIs: this can result in a 10^7^-fold increase in the product of SNR^2^ and the measurement speed (see Equations S82-S85 in (Szalay et al., 2016)). Therefore with each frame of the chessboard scanning method we only needed to record less than 5% of each VIP cell to preserve the fluorescence information for motion correction (Szalay et al., 2016). From these two parameters (and using Equations S82 and S84 from (Szalay et al., 2016)) we can calculate the increase in measurement speed and SNR for 3D chessboard scanning as follows:

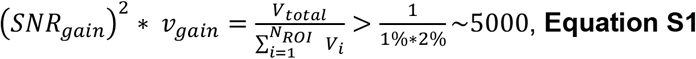

where *v_gain_* and *SNR_gain_* are the relative gains in measurement speed and SNR, respectively, *N_ROI_* is the total number of ROIs, *Vi* is the volume of region number *i*, and V_total_ is the total scanning volume. This means that we can get over 5000-fold increase in SNR^2^ or in the measurement speed (or even in the product of both) when AO-based 3D ROI scanning is used instead of point-by-point volume scanning.

To calculate this comparison more quantitatively, we compared 3D chessboard scanning with point-by-point scanning, volume scanning, and multi-layer imaging when AO scanning or resonant scanning with fast piezo z drive were used (see **Table S1**). We limited our comparison to these point scanning methods because they allow whole-field detection and, therefore, deep penetration in vivo. We recorded 120 VIP cells in a 689 μm × 639 μm × 580 μm volume with 548 × 507 × 193 pixel resolution using 3D chessboard scanning (**Figure 1A**). The 3D chessboard scanning method could image the 120 chessboards at 27.7 Hz (**Figure 1A**, **Table S1**). However, the measurement speed was only 0.00062 Hz when the same 120 neurons were recorded using point-by-point volume scanning when using the same, relatively long, pixel dwell time (30 μs). This means a 44762-fold lower measurement speed. We saw a smaller reduction in measurement speed when we compared chessboard scanning with resonant scanning. The highest speed of the currently available resonant scanners is about 16 kHz, corresponding to ~0.1 μs (=1 / 16 kHz / 548 pixel) pixel dwell time which results in a 0.16 Hz volume-scanning speed (**Table S1**) which is too slow to resolve Ca^2+^ responses. Moreover, as the pixel dwell time is 243-fold lower we would collect less signal from one pixel, resulting in a 41506-fold decrease in the product of SNR^2^ and measurement speed (**Table S1**). We could accelerate measurement speed by restricting the numbers of the recorded z layers to 19 because VIP neurons were present only in 19 z-layers in the exemplified measurement. However, in this case, we also needed to add about 20 ms setting time for each z layer because the long-range z drives required higher setting times according to the specifications of the piezo-actuators (see for example, https://www.physikinstrumente.com). This resulted in a measurement speed of ~1 Hz and in 6654-fold decrease in the product of SNR^2^ and measurement speed when compared to 3D chessboard scanning (**Table S1**). The increased SNR of the 3D chessboard scanning allowed the reduction of the laser intensity which resulted in lower phototoxicity according to the LOTOS (low-power temporal oversampling strategy (Chen et al., 2012)). The LOTOS-based multi-photon imaging is one of the main advantages of AO scanning and provides long lasting imaging in chronic behavior experiments.

During these comparisons we did not consider two important technical factors in our calculations. First: the gain in SNR was calculated only for a single pixel (which is a volume element in space, therefore we can name it as voxel). However, both 3D chessboard scanning and volume scanning capture multiple voxels from a single VIP neuron (in our measurements for chessboard scanning: 105.2 ± 0.4 voxels/neuron and for volume scanning: 338.5 ± 0.1 voxels/neuron). Therefore, in a more precise calculation we need to divide the improvement shown Table S1 for chessboard scanning with the ratio of 338/105.

Second: piezo actuators and resonant scanners are mechanically never perfectly balanced and are also sensitive to local mechanical vibrations and thermal turbulences which results in tumbling, wobbling, and jitter in the laser scans. These mechanical effects are difficult to precisely quantify into fluorescence changes although they would compensate the first factor. Therefore, for simplicity of calculation, both factors were ignored in our calculations.

Random-access targeting of regions of interest by AO scanning is useful not only in 3D but also in two-dimension (2D). Because the ratio of the VIP cells in the cortex is about < 1% we can estimate the increase in measurement speed and SNR in 2D as follows:

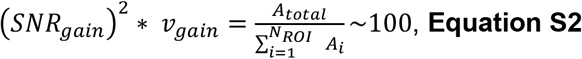

### Visual stimulation

An LCD monitor was placed at a distance of 20 cm from the contralateral eye of the mouse, spanning 100° x 70° of the visual field. The objective was covered with a black rubber shield to prevent stray light entering through the gap between the animal’s head and the objective. A visual stimuli protocol written in Matlab using the ‘Psychtoolbox’ package. The protocol consisted of eight differentially directed gratings with an angular interval of 45°. At the beginning of each trial, a gray screen was presented for 20s. After that, a grating appeared and remained still for 1 s, and then moved orthogonally to its orientation for 6 s at 1 cyc/s speed, then it stayed still for 1 s, and finally the grey screen reappeared again. Gratings were repeated 10-20 times per direction in pseudorandom order. Pyramidal cell data were obtained from Thy-1-Cre mice.

### Two-photon imaging data analysis

Motion correction, selection of ROIs corresponding to VIP cells on the frames of 3D chessboard scanning, background calculation, ΔF/F calculation, filtering and data visualization was performed using the MES data acquisition software written in Matlab and C++ (Femtonics). Motion correction, if necessary, was conducted with a custom-written offline motion correction algorithm (see Off-line motion corrections section), and remaining artefacts were interpolated or smoothed with partial Gauss filtering under visual control.

For the trial-to-trial analysis we considered a neuron in a given trial as active if the difference between the peak ΔF/F value of reinforcer delivery epoch (0-2 sec interval after reinforcement onset) and mean ΔF/F value of baseline epoch (−2-0 sec interval before tone onset) was higher than 2 standard deviations (SD). Peak ΔF/F value was defined here as the average ΔF/F value of the datapoints around the peak in the range of 250 ms. The results presented in **Figure S2** used two SD as a cutoff. We defined synchronicity as the number of active neurons divided by the total number of all neurons in a given a trial, i.e. how many neurons are activated simultaneously. Reliability was calculated as the number of active trials divided by the total number of trials, i.e. how reliably the neuron is activated in Hit and FA trials.

We used linear regression models to address heterogeneity of the cue responses (**Figures 2A, S3D**). The explanatory variable for the first model was the hit rate (number of hit trials divided by the number of go trials) to characterize behavior. The dependent variable was the relative size of the average cue response compared to the average reinforcement response as a reference. The values were calculated on the population average traces of the hit trials from each measurement. In the second model we used categorical variables with dummy coding for 4 functional regions (motor, sensory, parietal, visual) as explanatory variables to describe regional differences. The regression was fitted using ordinary least squares method of Statsmodels package in Python 3 based Anaconda data science platform.

The clustering analysis (**Figures 2F-G, S3A-B**) was done using a custom Matlab routine. Positive somatic Ca^2+^ responses recorded during hit trials were extracted. Data were z-scored using the mean and variance of fluorescence during the first second of recording. We furthered normalized using the maximal amplitude of the response calculated during the period from 0 to 4s after reward delivery. We applied a dimensionality reduction along the time axis using a principal component analysis on the period from 0 to 4s after reward delivery. We then considered only the first 4 principal components (PCs) explaining 90% of the data for clustering purposes. K-means clustering with 5 replicates was used to cluster into 5 types the PCs of the responses of VIP interneurons.

We applied Tensor Component Analysis (TCA) (Kolda and Bader, 2009; Williams et al., 2018) on somatic Ca^2+^ responses recorded during the discrimination task. After smoothing, single trial neural activities corresponding to the reaction time periods for hit and FA trials were time-wrapped to a fixed 1.5 sec / 30 data points in length. All recordings were rendered non-negative by subtracting the minimal fluorescent DF/F value for each cells. Data were finally normalized by dividing by the average maximum fluorescent ΔF/F value of Hit-only trials. Only the first 10 trials of each types were then selected for each cell and each session and assembled in a NxTxK matrice where N=771 neurons, T: time (s), K=40 trials. TCA reconstruction error was computed with different latent numbers [1,2,3,4,5,10,15,20] with 10 different initial conditions for each latent number. Using 3 latents lead to a reconstruction error of 21%.

### Pupil diameter

The video recorded with the camera was first thresholded to isolate the pupil on the image. The pupil area was fitted to an ellipse, and the main diagonal was extracted. Missing frames caused by spontaneous or air puff-triggered blinking were interpolated manually. A Gaussian filter was applied to smooth eye movement-related artefacts. In analyzing the change of pupil diameter, the traces were normalized to ΔP/P = (P(t)-P_0_)/P_0_ using a two second period before tone onset as baseline (P0). Trials of each outcome were separated to high- and low-arousal groups on the basis of the change in the pupil diameter. The area under the pupil diameter curve was calculated in the 0-3 sec interval after reinforcement onset, and the median value was selected. If the area under the curve value of a given trial was higher or lower than the median value, it was considered to be high- or low-arousal trial, respectively. For baseline pupil analysis, when we compared the VIP cell activity after reinforcement associated with low and high baseline arousal levels, the trials were again separated into low- and high-arousal trials, but here, the basis of the separation was the area under the pupil diameter curves in the [−2;0] sec interval before the tone onset.

### Locomotion velocity analysis

Velocity traces were first Gauss filtered. In Hit and FA trials, we defined the change in the running speed as the speed difference between the reinforcer delivery time period (0-2 sec interval after reinforcement onset) and baseline time period (−2-0 sec interval before tone onset). In Miss and CR trials, the speed difference was calculated between the tone delivery time period (−0-2 sec interval after tone onset) and baseline time period (-2-0 sec interval before tone onset). Trials were separated into low and high speed change groups according to the median speed change value.

### Visual stimulation

Orientation and direction selectivity indexes were calculated as OSI = (R_pref_ - R_ortho_)/(R_pref_ + R_ortho_), and DSI = (R_pref_ - R_opp_)/(R_pref_ + R_opp_) (Schumacher et al., 2019), where R_pref_ denotes the amplitude of the response to the preferred orientation (OSI) or direction (DSI), R_ortho_ denotes the response to the orthogonal orientation in the OSI formula, and Ropp refers to the response to the direction that is opposite to the preferred one.

### Statistical analysis

For all analyses, the activation/suppression period was set to be 0-2 sec after stimulus onset: the statistical significance of the change of Ca^2+^ responses was then evaluated and compared to a stimulus-free baseline (−2-0 sec before stimulus onset). The statistical significance of the activation and suppression was determined by a P value cutoff of 0.05. First, the mean baseline values of each trial were subtracted from each Ca^2+^ trace in order to directly compare the effect of stimuli on Ca^2+^ responses and minimize the effect of unknown sources of noise. Lilliefors normality test was used to evaluate whether the Ca^2+^ signals of individual VIP neurons followed a normal distribution. The Lilliefors test showed that 76% of VIP neurons (Hit: 70%, FA: 82%) followed a normal distribution. The fraction of the neurons activated by reinforcers (see below) with normal distribution (91%) was similar to that of activated neurons with non-normal distribution. Therefore, we used a one-tailed one sample t-test to classify the activation and suppression (see Supplemental Information for a more sensitive analytical method). Neurons were classified as responsive when either activation or suppression was statistically significant. Student’s t-test (\**p* < 0.05, \*\**p* < 0.01, \*\*\**p* < 0.001) was also used to compare calcium responses associated with low and high arousal, or low and high running speed. PCA loadings of different areas and TCA trial factors of trialtypes with and w/o reinforcement were compared with Mann-Whitney test. If not otherwise indicated, data are presented as mean ± SEM.

### Off-line motion correction

In the case of chessboard scanning, neuronal somata were selected from a z stack, then the selected square regions of interest were arranged as a 2D chessboard. Since the motion of a single frame during the scanning period as well as the relative rotation between subsequent frames, were not relevant, the transformation to be corrected could be approximated by a simple translational transformation between the scanning periods of the different frames. For efficiency, an algorithm based on fast Fourier transformation was used (Fuster and Bressler, 2015; Guizar-Sicairos et al., 2008). The template images were chosen either manually or by selecting the best correlating 20% of the relevant ROIs on all frames.

In some cases, images were also preprocessed, by either adaptive histogram equalization or simple median filtering. The image registration algorithm also provided the error of matching the moving images to the template images. As the drifting and scanning parameters were identical for each scanned ROI, we calculated the final displacement vector as the median of a fixed percentage of all ROIs with the smallest matching errors.

## MULTIMEDIA FILES

**Movie S1. related to Figure 1. Recording sparse interneuronal population in large volume**

Z-stack from half mm^3^ neocortical volume was obtained in the parietal cortex. Then small squares containing the VIP interneurons’ somata were selected as ROIs. The squares were rearranged to form a 2D matrix to track the cell activity.

**Movie S2. related to Figure 1. VIP population activity during an auditory discrimination task.**

Example false alarm trial of an imaging session with pupillometry, velocity recording and motion corrected calcium imaging of 52 VIP interneurons. Flashing white speaker and red air cloud icons mark the tone and air puff onsets.

## DATA AVAILABILITY

Data are available upon request from Balázs Rózsa and Adam Kepecs.

## CODE AVAILABILITY

Custom written analysis codes are available at https://github.com/QuentinNeuro.

## AUTHOR CONTRIBUTIONS

The project was initiated and the experiments were conceived by A.K, H.P., B.R., Z.S., carried out by Z.S., H.P., Q.C., B.C, and analyzed by Z.S., H.P., Q.C., K.Ó. TCA was performed by K.Ó. and Q.C. Scanning strategies were developed by G.K. and K.Ó. F.A. helped with fiber photometry. Z.S., H.P., Q.C., B.R. and A.K. wrote the manuscript with comments from all authors.

## ACKNOWLEDGEMENTS

We thank Lídia Popara, Áron Szepesi, and Alexandra Bojdán for technical help, and all members of the Kepecs and Rózsa labs for their helpful comments. This study was supported by KFI-2018-00097, VKE-2018-00032, NKP-2017-00001, KTIA_NAP_12-2-2015-0006, 2017-1.2.1-NKP-2017-00002, GINOP_2.1.1-15-2016-00979, and János Bolyai Research Scholarship of the Hungarian Academy of Sciences. The project was implemented with the support from the National Research, Development and Innovation Fund of Hungary financed under the VKSZ_14-1-2015-0155, KFI_16-1-2016-0177, NVKP_16-1-2016-0043 funding scheme. This project received funding from the European Research Council (ERC) under the European Union’s Horizon 2020 research and innovation programme (grant agreement No. 682426 and VISGEN_734862, 712821-NEURAM). H.P. is supported by NARSAD Young Investigator Grant and NIH R01MH110391. The study received funding from the National Institutes of Health, R01NS075531 and R01NS088661, to A.K.

## DECLARATION OF INTERESTS

G.K. and B.R. are founders of Femtonics Ltd. B.R. is a member of its scientific advisory board.

## SUPPLEMENTARY INFORMATION

**Figure S1. related to Figure 1.**
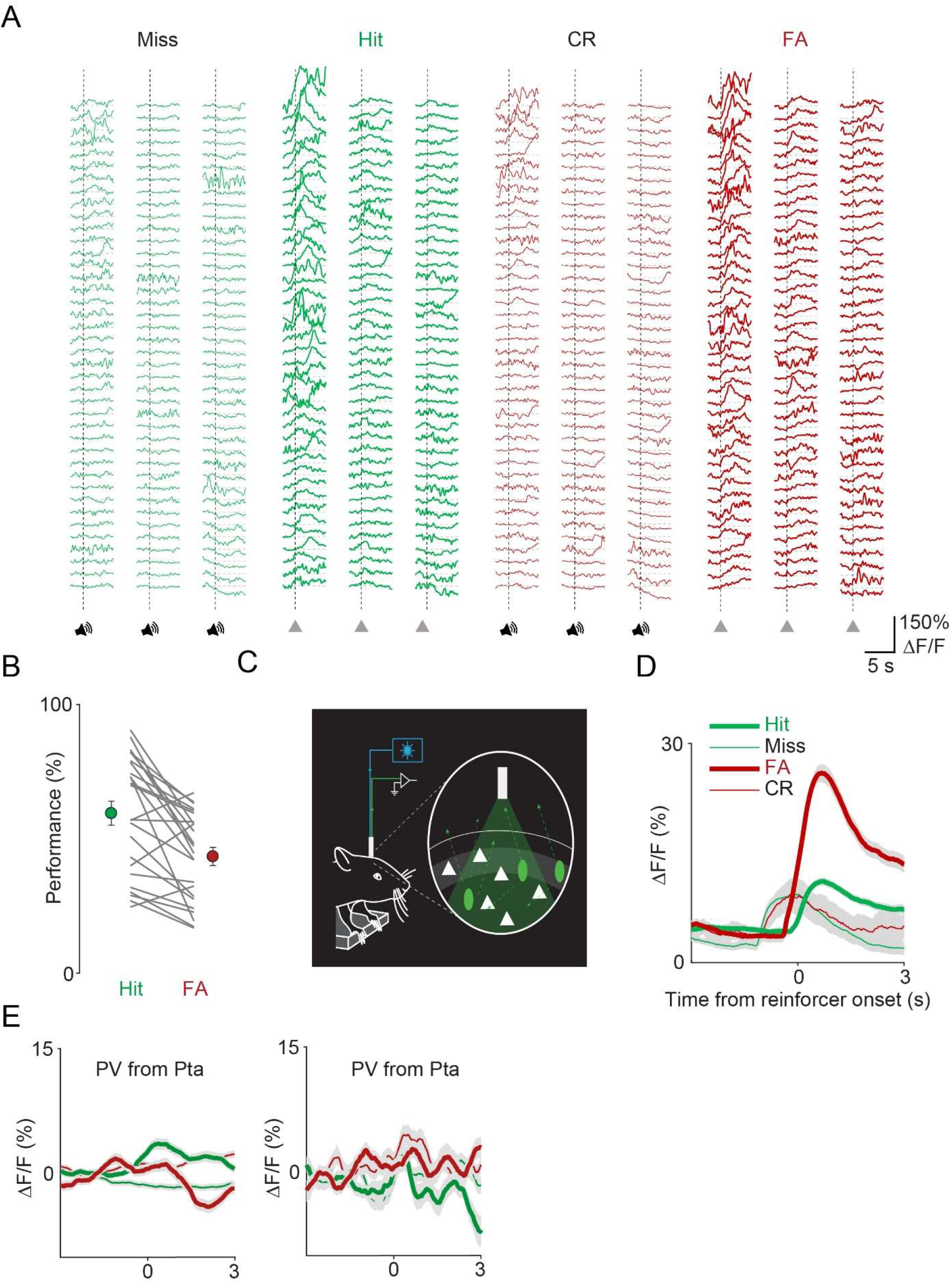
3D-random-access two-photon imaging and fiber photometry of VIP neurons in an auditory discrimination task. **A)** The somatic Ca^2+^ responses in **Figure 1C** shown in transient form **B)** Hit and FA rate of the mice during the imaging and fiber photometry sessions (n=24 sessions, n=22 mice). **C)** Schematics of fiber photometry experiments. **D)** Average transients of VIP interneurons (mean±SEM) for Hit (thick green), FA (thick red), Miss (thin green) and CR (thin red) recorded from the ACx using fiber photometry. **E)** Average transients of PV interneurons (mean±SEM of n=17 and n=37 cells) for Hit (thick green), FA (thick red), Miss (thin green) and CR (thin red) recorded from the parietal cortex.

**Figure S2. related to Figure 2.**
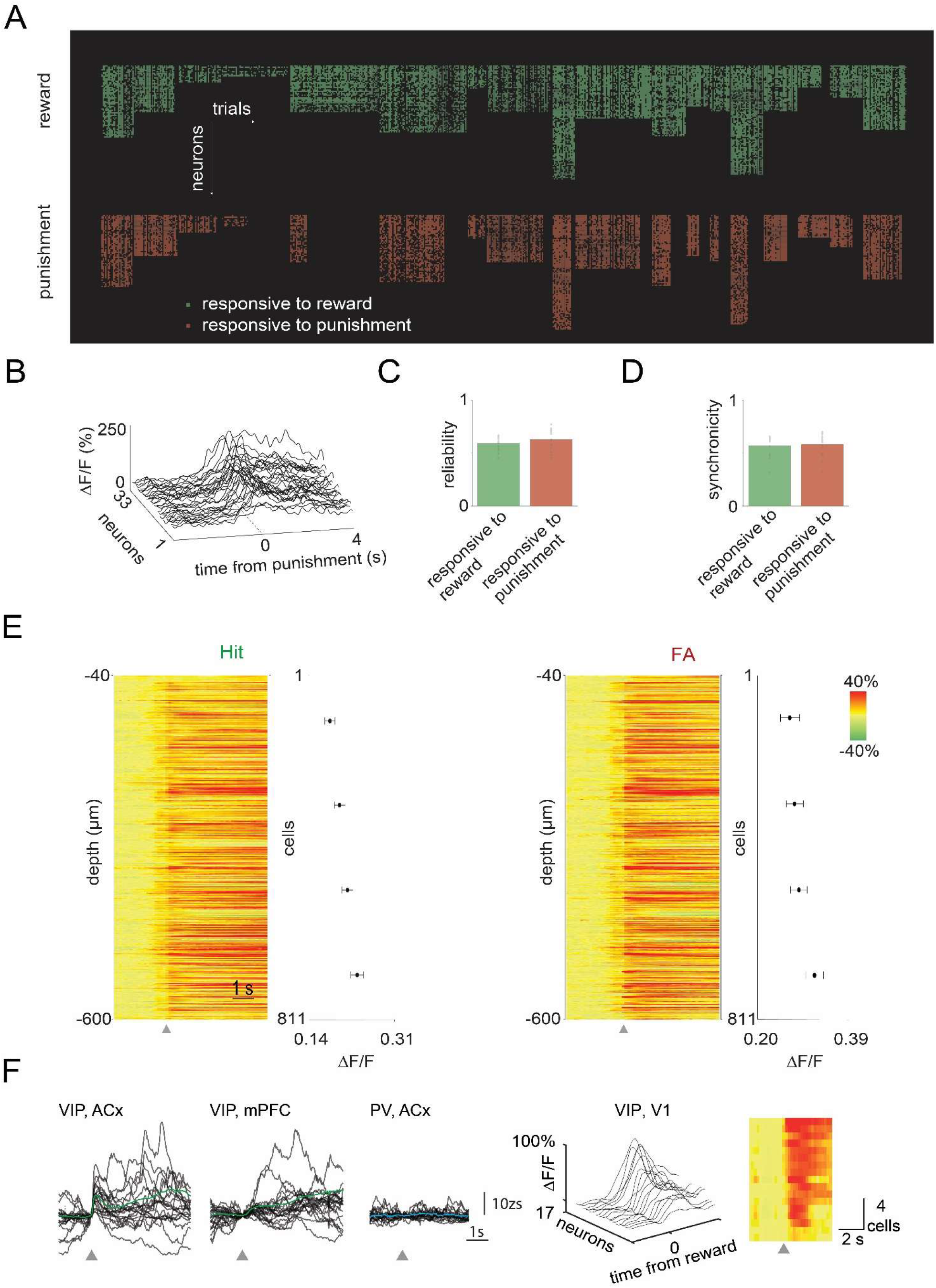
Quantification of the activity of VIP neurons across the dorsal cortex. **A)** Raster plot of the trial-to-trial activation of the responsive VIP neurons in Hit and FA trials during the two-photon imaging sessions (n=18 sessions, n=746 cells). **B)** An example of the synchronous activation of the VIP neurons in a FA trial. **C)** Reliability of the VIP neurons in Hit and FA trials. **D)** Synchronicity of the VIP neurons in Hit and FA trials. **E)** Left, raster plot of the average responses of the VIP neurons in Hit trials ordered according to their cortical depth. Graph shows the binned maximums of the averaged responses (bin size = 223 cells). Right, raster plot and graph for FA trials. Gray triangles mark reinforcement onset. **F)** Single trial (black) and average (green or blue) VIP or PV interneuron z – scored activity recorded using fiber photometry during uncued reward delivery. Grey triangle shows the onset of the uncued reward. Right, single trial and average activity of VIP interneurons from V1 recorded with 2 – photon microscope during uncued reward delivery.

**Figure S3. related to Figure 2.**
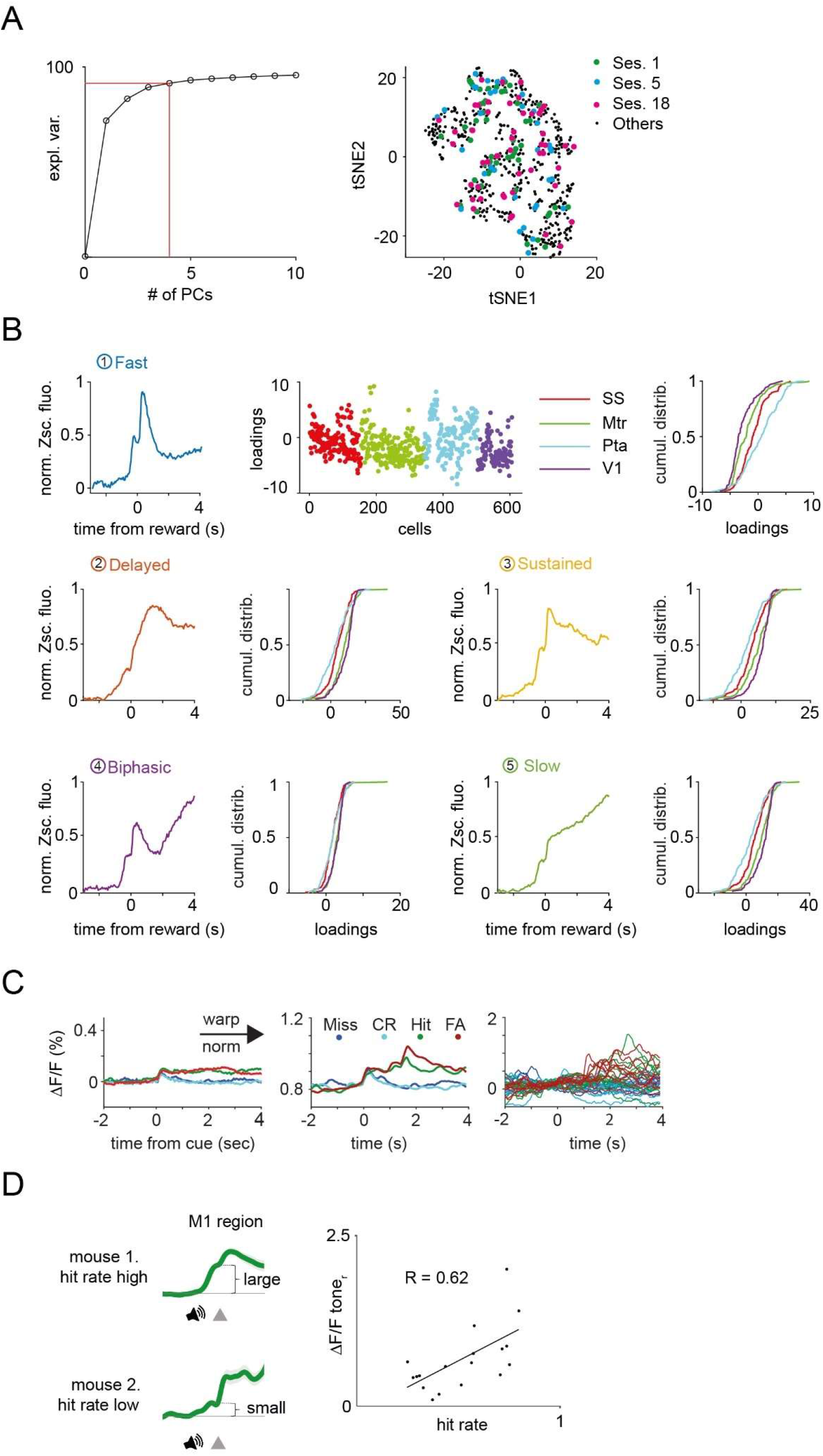
Heterogeneity in VIP neuronal responses across the dorsal cortex. **A)** Left, explained variance. We used 5 PCs, explaining >90% of the variance of our data for the k-means clustering. Right, neurons from individual recording sessions are scattered in the tSNE space. **B)** Top, the first principal component, corresponding loadings, and regional cumulative distributions of the loadings. Middle and bottom, remaining principal components and the regional cumulative distributions of the corresponding loadings. **C)** Average activity of 17 VIP interneurons for different trial types before (left) and after (middle) TCA preprocessing. After smoothing, single trial neural activities corresponding to reaction time periods for hit and FA trials were time-wrapped to a fixed 1.5 sec in length. All recordings were rendered non-negative by subtracting the minimal ΔF/F value for each cell. Data were finally normalized. Right, only the first 10 trials of each type were selected for the TCA. **D)** Heterogeneity of the cue responses. Left, higher hit rate was associated with larger tone related response components in the population average traces of Hit trials. Left, scatter plot of the size of the average tone response and the hit rate.

**Figure S4. related to Figure 3.**
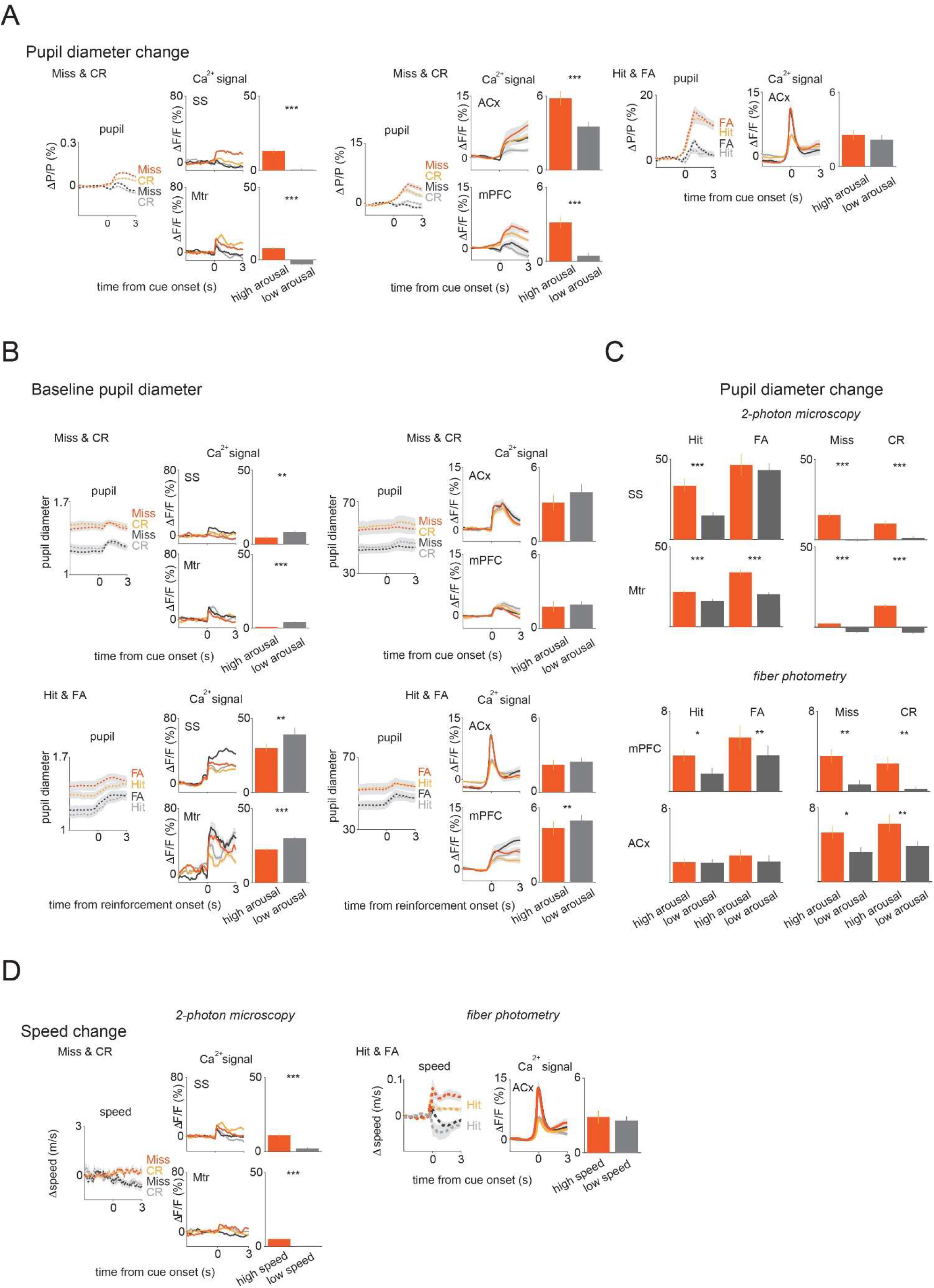
The baseline and the change in pupil diameter, and the change of speed additionally modulate VIP neuronal activity on top of activation by cues and outcomes. **A)** Population averages for Miss and CR (left and middle) during high and low arousal change in the SS and Mtr regions (left), and ACx and mPFC regions (middle). Right, population averages for Hit and FA in ACx. Bars indicate average peak amplitudes (mean±SEM). **B)** Population averages for Miss and CR (top) and Hit and FA (bottom) by high and low baseline arousal levels in the SS and Mtr regions (left), and ACx and mPFC regions (right). Bars indicate average peak amplitudes (mean±SEM). **C)** Average peak amplitude bars for Hit, FA, Miss and CR separately during high and low pupil change in the SS, Mtr, mPFC and ACx regions. (mean±SEM). **D)** Population averages for Miss and CR (left) during high and low speed change in the SS and Mtr regions (left) and for Hit and FA in ACx (right). Bars indicate average peak amplitudes (mean±SEM).

**Figure S5. related to Figure 4.**
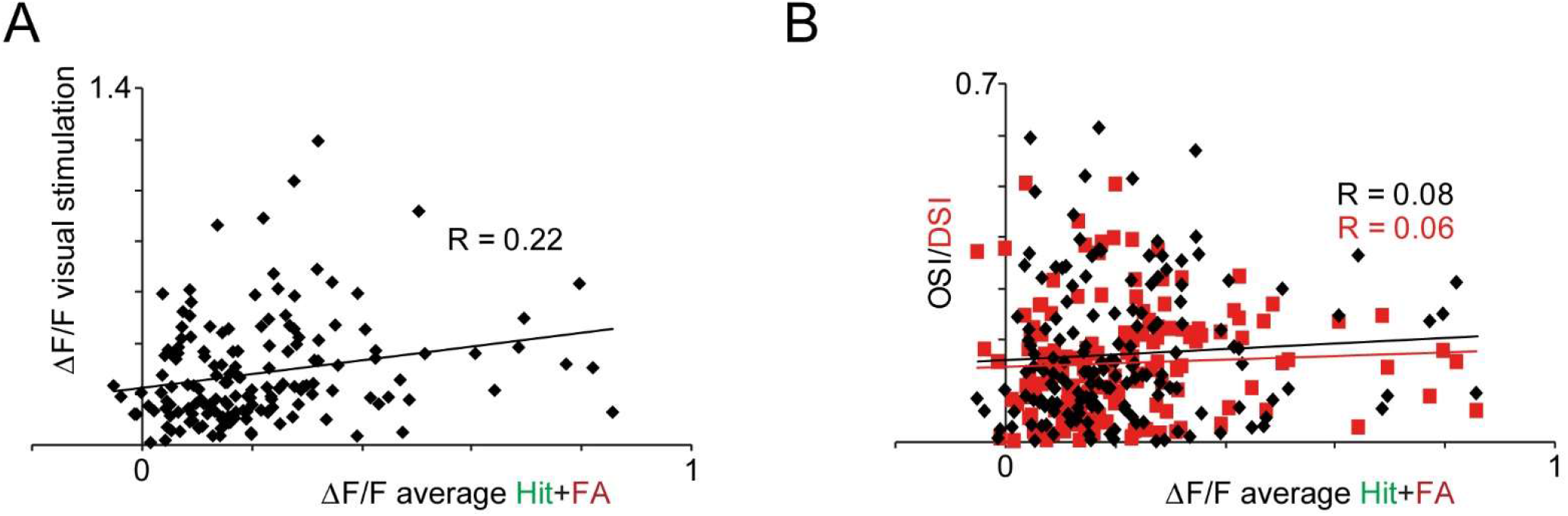
Quantification of the connection of the visual tuning parameters and reinforcement-related responses. **A)** Scatter plot of reinforcement-vs visual stimulation-induced responses of the same VIP cells. **B)** Scatter plot of reinforcement-induced responses vs. OSI or DSI parameters of the same VIP cells.

**Table S1. related to Figure 1.**
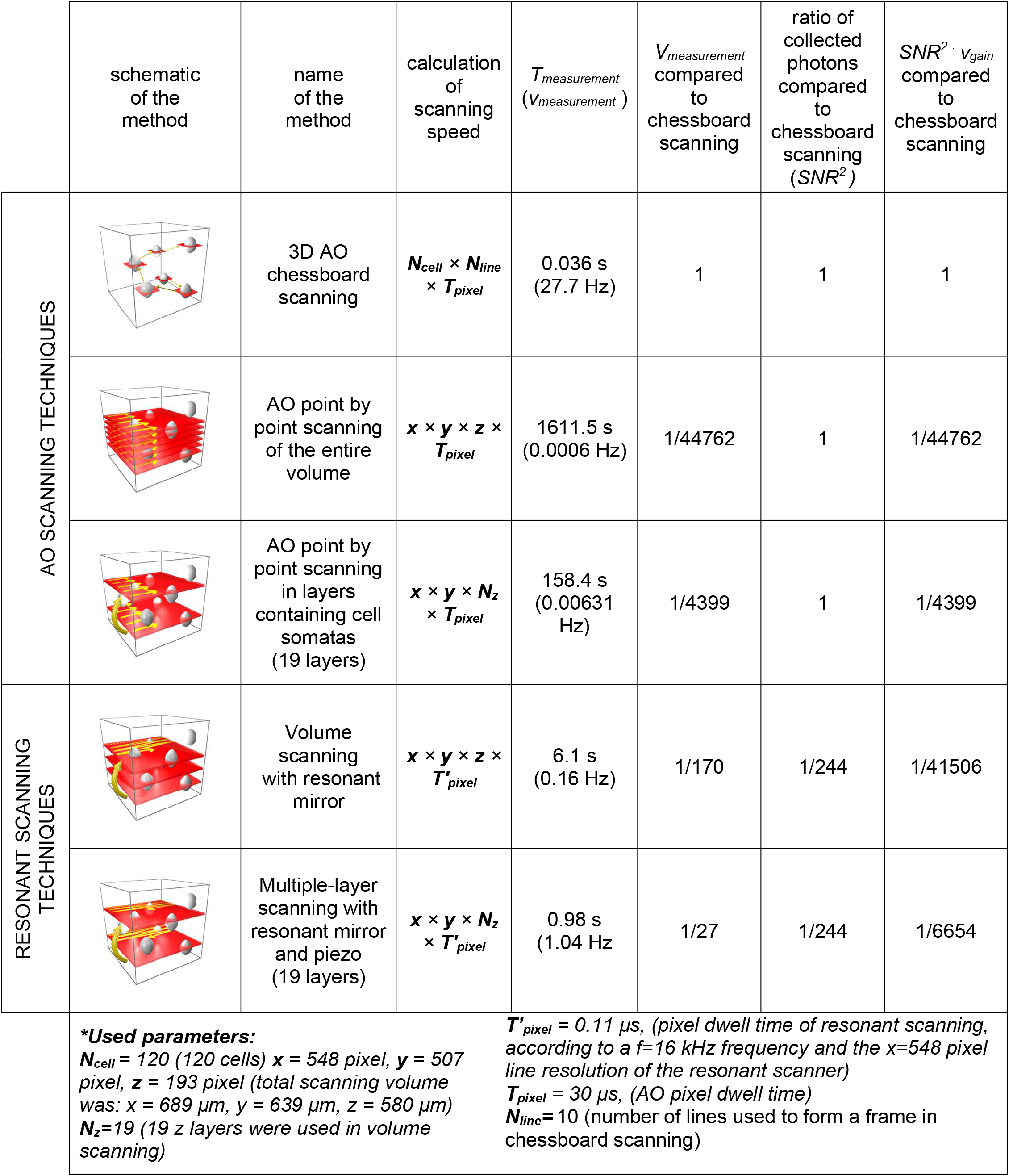
Comparison of different scanning methods.

| | schematic of the method | name of the method | calculation of scanning speed | $T_{\text{measurement}}$ ($V_{\text{measurement}}$) | $V_{\text{measurement}}$ compared to chessboard scanning | ratio of collected photons compared to chessboard scanning ($SNR^2$) | $SNR^2 \cdot V_{\text{gain}}$ compared to chessboard scanning |
| --- | --- | --- | --- | --- | --- | --- | --- |
| AO SCANNING TECHNIQUES | | 3D AO chessboard scanning | $N_{\text{cell}} \times N_{\text{line}} \times T_{\text{pixel}}$ | 0.036 s (27.7 Hz) | 1 | 1 | 1 |
| | | AO point by point scanning of the entire volume | $x \times y \times z \times T_{\text{pixel}}$ | 1611.5 s (0.0006 Hz) | 1/44762 | 1 | 1/44762 |
| | | AO point by point scanning in layers containing cell somatas (19 layers) | $x \times y \times N_z \times T_{\text{pixel}}$ | 158.4 s (0.00631 Hz) | 1/4399 | 1 | 1/4399 |
| RESONANT SCANNING TECHNIQUES | | Volume scanning with resonant mirror | $x \times y \times z \times T'_{\text{pixel}}$ | 6.1 s (0.16 Hz) | 1/170 | 1/244 | 1/41506 |
| | | Multiple-layer scanning with resonant mirror and piezo (19 layers) | $x \times y \times N_z \times T'_{\text{pixel}}$ | 0.98 s (1.04 Hz) | 1/27 | 1/244 | 1/6654 |
| **\*Used parameters:**<br>$N_{\text{cell}} = 120$ (120 cells) $x = 548$ pixel, $y = 507$ pixel, $z = 193$ pixel (total scanning volume was: $x = 689 \mu\text{m}$, $y = 639 \mu\text{m}$, $z = 580 \mu\text{m}$)<br>$N_z=19$ (19 z layers were used in volume scanning) | | | | $T'_{\text{pixel}} = 0.11 \mu\text{s}$, (pixel dwell time of resonant scanning, according to a $f=16$ kHz frequency and the $x=548$ pixel line resolution of the resonant scanner)<br>$T_{\text{pixel}} = 30 \mu\text{s}$, (AO pixel dwell time)<br>$N_{\text{line}}=10$ (number of lines used to form a frame in chessboard scanning) | | | |

Scanning speed was calculated according to the equations in the column “calculation of scanning speed”. Ratio of collected photons was calculated from relative pixel dwell times. All parameters used for calculations are listed in the bottom field. Note, that chessboard scanning provides 170-fold faster measurement speed and 244-fold higher photon collection compared to volume scanning with resonant mirrors.

**Table S2.** Calculation of ratio of responsive neurons following reward.

| Animals | #1 | #2 | #3 | #4 |
| --- | --- | --- | --- | --- |
| total number of cells recorded | 40 | 40 | 39 | 34 |
| number of cells with low SNR | 4 | 1 | 0 | 2 |
| # of cells with low SNR | 1,9,19,20 | 12 | 0 | 9,12 |
| number of non-responsive cells | 4 | 1 | 1 | 1 |
| # of cells with no response (#) | 3,10,28,32 | 30 | 30 | 5 |
| responsive cells | 36 | 39 | 39 | 32 |
| Ratio of responsive cells (%) | 88.88 | 97.44 | 97.44 | 96.88 |
| Average ratio (% , mean±SEM) | 95.16±2.09% |  |  |  |

### Baseline arousal level modulates VIP activity

We split the trials based on the baseline or inter-trial arousal level (**Figure S4B**). Thus, trials were split by the median value of the baseline pupil diameter. This analysis revealed that the arousal level in the baseline period inversely correlated with the degree of VIP activation by reinforcers. For instance, when baseline arousal level was low, the reinforcers tended to induce a stronger increase in VIP activity. When baseline arousal level was high, the reinforcers induced a smaller increase (high vs. low baseline: SS: 30% vs 39% (ΔF/F), n=26, p<0.01; Mtr: 23% vs 30% (ΔF/F), *n*=111, *p*<0.001; **Figure S4B**). Interestingly, the anti-correlation was also observed between baseline pupil diameter and the increase in pupil diameter. When the baseline pupil diameter was small, the increase in pupil diameter tended to be higher and vice versa (**Figure S4B**). Taken together, these results indicate that the baseline arousal level is another important factor that modulates VIP activity.

#### Identifying responsive neurons

In this section, we will introduce a new method for selecting responsive neurons from large neuronal populations recorded simultaneously. The method is based on the following: 1) a thresholding method in which neurons with very poor SNR are eliminated at the beginning of the analysis; 2) baseline calculation in the pre-stimulus period; and 3) one sample t-test in the response period.

##### 1) The thresholding method

Majority of the cells showed robust spontaneous and reinforcement-related responses with variable amplitude and frequency. However, in some neurons, responses were below the detection threshold in a recording period of over 5 minutes. To eliminate neurons with low SNR we calculated the mean amplitude of the 15±5% largest peaks detected in Ca^2+^ transients during the recording period of over 5 minutes and divided it by the average, pre-stimulus STD. Neurons with a (mean amplitude)/STD ratio below 5 were eliminated from the analysis. This threshold eliminated 4.98±0.01% of the VIP cells (**Table S2**).

##### 2) Baseline calculation

There are many ways to define a neuron as responsive or non-responsive. For all definitions we need a baseline relative to which responsiveness can be calculated. The simplest approach is to define a pre-stimulus temporal interval ([T_01_,T_02_]) before the cue onset as a baseline period, and calculate the mean, μ_0_ (see below).

##### 3) One-sample t-test

In a one sample t-test, the null hypothesis is that the population mean is equal to a specified value (μ0).

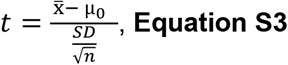

where SD and n are the standard deviation and sample size, respectively. The degrees of freedom (DF) is n-1. The distribution of the population of sample means 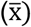 is assumed to be normal although this is not required for the parent population. The distribution of t_p_ will be approximately normal N(0,1) according to the central limit theorem. If we substitute SD with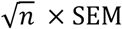 where SEM is the standard error of the mean, we get the following criterium:

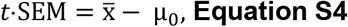

In Student’s t-test the 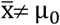 hypothesis is accepted as significant if |*t*| > *t_p_* where *t_p_* is defined as (*P*(|*t*| > *t_p_*) = *p*, where P denotes probability. The Student’s distribution defines *t_p_* at a given DF *(n-1)* and a given *p* value. Therefore, we can define the following criterium for significance:

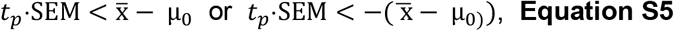

The simplest approach to define a neuron as responsive is to calculate in a one-sample t-test whether the mean response 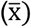 of the neuron is significantly larger (or smaller) in a given interval after 0 ms (where 0 ms is the time of the stimulus) than the baseline average value (μ_0_, which is equal to zero by definition). According to this definition and **Equation S5**, a neuron is responsive if its average time-dependent response transient, 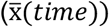 is larger during a given time interval than the product of *t_p_* and the SEM of the population:

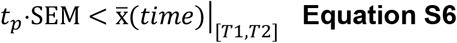

and for significantly smaller responses (for inhibition) we can use the modified second Equation from **Equation S5**:

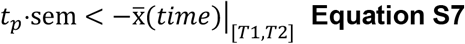

In practice, we defined the pre-stimulus baseline period, from 2 s before cue onset to the cue onset time. The interval of responses was defined from the time of the reinforcement time (0 s) to 2 s after the reinforcement. This also means that both *T_1_* and *T_2_* time values must be part of the [0 s, 2 s] response interval. In theory, there is no limit to the minimum length of the [T_1_, T_2_] interval; however, in practice we used the T_2_-T_1_ ≥ 500 ms criterium which was in the range of the mean length of the single AP potential-induced response at half maximum.

The number of reward and punishment transients collected in a given experiment was variable (between 30 and 50) resulting in variable *t_p_* values which had to be calculated for each experiment separately. For example, a trial with 34 transients means 33 degrees of freedom (34-1) and *t_p_* = 1.692 (*p*<0.05; two tails). We performed fast 3D recording of VIP interneurons (34-40 cells, from 4 mice, **Table S2**) and calculated the activation ratio. The activation ratio for reward was 95.16±2.09%, which is a much higher ratio than that determined using the standard one sample t-test above (76.22±9.65%).

